# Dipeptides are minimalistic but sufficient for liquid-liquid phase separation

**DOI:** 10.1101/2021.04.03.438298

**Authors:** Yiming Tang, Santu Bera, Yifei Yao, Jiyuan Zeng, Zenghui Lao, Xuewei Dong, Ehud Gazit, Guanghong Wei

## Abstract

Liquid-liquid phase separation (LLPS) of proteins mediates the assembly of biomolecular condensates involved in physiological and pathological processes. Identifying the minimalistic building blocks and the sequence determinant of protein phase separation is of urgent importance but remains challenging due to the enormous sequence space and difficulties of existing methodologies in characterizing the phase behavior of ultrashort peptides. Here we demonstrate computational tools to efficiently quantify the microscopic fluidity and density of liquid-condensates/solid-aggregates and the temperature-dependent phase diagram of peptides. Utilizing our approaches, we comprehensively predict the LLPS abilities of all 400 dipeptide combinations of coded amino acids based on 492 micro-second molecular dynamics simulations, and observe the occurrences of spontaneous LLPS. We identify 54 dipeptides that form solid-like aggregates and three categories of dipeptides with high LLPS propensity. Our predictions are validated by turbidity assays and differential interference contrast (DIC) microscopy on four representative dipeptides (WW, QW, GF, and VI). Phase coexistence diagrams are constructed to explore the temperature dependence of LLPS. Our results reveal that aromatic moieties are crucial for a dipeptide to undergo LLPS, and hydrophobic and polar components are indispensable. We demonstrate for the first time that dipeptides, minimal but complete, possess multivalent interactions sufficient for LLPS, suggesting that LLPS is a general property of peptides/proteins, independent of their sequence length. This study provides a computational and experimental approach to the prediction and characterization of the phase behavior of minimalistic peptides, and will be helpful for understanding the sequence-dependence and molecular mechanism of protein phase separation.

**Significance:** Protein liquid-liquid phase separation (LLPS) is associated with human health and diseases. Identifying the minimalistic building blocks and sequence determinants of LLPS is of urgent importance but remains computationally challenging partially due to the lack of methodologies characterizing the liquid condensates. Herein we provide approaches to evaluate LLPS ability of dipeptides, and screen all 400 dipeptides by MD simulations combined with multi-bead-per-residue models which capture key interactions driving LLPS that are missing in one-bead-per-residue models. Three categories of LLPS dipeptides are identified and the experimentally-verified QW dipeptide is by far the smallest LLPS system. Our results suggest that dipeptides, minimal but complete, possess multivalent interactions sufficient for LLPS, and LLPS is a general property of peptides/proteins, independent of their length.

## Introduction

Accumulating evidence demonstrates the importance of protein phase separation in physiological and pathological processes (1–3). Liquid-liquid phase separation (LLPS) of proteins underlies the formation of membraneless compartments in cells (4, 5), possessing important intercellular functions, such as facilitating bio-chemical reactions, activating stress response, and regulating gene transcriptions (6–8). In addition, aberrant LLPS and the liquid-solid phase separation (LSPS) can lead to formation of neurotoxic aggregates such as amyloid plaques in Alzheimer’s disease, fibrillary aggregates in Amyotrophic Lateral Sclerosis (ALS) (9, 10), aberrant stress granules in Frontotemporal Dementia (FTD) (11), and Lewy bodies in Parkinson’s disease (12). Identifying the sequence determinants and the molecular mechanisms underlying phase separation is yet an unmet need crucial for designing therapeutic targets to treat protein-aggregation disorders.

A rapidly increasing number of proteins have been demonstrated to have LLPS capabilities. Examples include the transactive response DNA binding protein (TDP-43) (13, 14), Tau (15–17) and Fused in Sarcoma (FUS) (18–20). It has been reported that LLPS is driven by multivalent interactions entailed by the amino acid sequence (1, 21, 22). In addition, it is considered that a minimum sequence length of protein is required and below which LLPS cannot occur. For example, the LLPS of the poly-PR/poly-GA requires a minimum peptide length of 50/100 (23, 24). In contrast, numerous experiments demonstrate that even very short peptides can self-assemble into ordered solid-like nanoarchitectures. For example, FF, FW, and IF can respectively form nanotubes, tubular and fibrous nanostructures (25–27). In this context, we suspect that ultrashort peptides, as short as dipeptides, may also possess LLPS capacity. In order to identify the minimalistic building blocks and the sequence determinant of protein phase separation, we decipher the LLPS/LSPS ability of all 400 dipeptide combinations of coded amino acids by conducting a combined computational and experimental study.

Different theoretical and computational approaches have shown success in probing the phase behaviors of intrinsically disordered proteins (IDPs) or proteins with intrinsically disordered regions (IDRs) (28, 29). Such methods include random phase approximation theory by Lin and Chan (30), field-theoretic simulations by Shea *et al*. (31), Gibbs-ensemble simulations by Zhou *et al*. (32, 33), and slab simulations by Mittal *et al*. (34–36) and others (37). Those studies greatly enhance our understanding of key physical principles and sequence features crucial to phase separation of IDPs or IDR-containing proteins. Following these studies, we have attempted to investigate the phase separation of dipeptides by utilizing the slab simulation protocol in combination with the hydrophobicity-scaled (HPS) one-bead-per-residue model. Unexpectedly, dipeptides with known high aggregation capability (such as FF, FW and IF) display totally no sign of aggregation (Supplementary Part 1), probably due to the fact that highly-simplified protein models significantly reduces the molecular interaction sites, thus lack molecular details to explicitly describe aromatic stacking, hydrogen-bonding interactions and peptide-water interactions crucial for ultrashort peptide phase separation (22, 38–40). On the other hand, the slab simulation protocol, which has become a *de facto* standard for computing the LLPS phase diagrams (29), underestimates the density of dilute phases (i.e., the saturation concentration for phase separation) probably due to the lack of explicit solvents. Thus, a more detailed description of ultrashort peptides in combination with explicit solvent model is necessary for the investigation of their LLPS.

Hereby, we have carried out micro-second explicit solvent molecular dynamics (MD) simulations on each of the 400 dipeptides in combination with the multiple-bead-per-residue MARTINI model which explicitly capture specific physical interactions (such as aromatic stacking, and protein-water interactions). A similar simulation protocol has been used by Tuttle and coworkers to screen out dipeptides with high self-assembly ability (41), albeit with a short simulation time scale. Nevertheless, the LLPS ability of dipeptides and their phase behaviors remain completely elusive. We have developed a novel, efficient approach to predict the LLPS propensities of ultrashort peptides which consists of two steps: (1) the aggregation capability of each dipeptide is quantified by the clustering degree and the collapse degree of initially randomly dispersed dipeptide molecules, and (2) we define two physical parameters (the fluctuation extent of clustering degree and the exchange rate of inter-peptide interactions) to characterize the microscopic fluidity of the aggregates and distinguish between LLPS and LSPS. Our extensive simulations demonstrate that dipeptides can spontaneously phase separate into solid- or liquid-like condensates, and predict the LLPS and LSPS propensities of all 400 dipeptides, among which QW possesses the highest LLPS propensity. Our predictions are further verified through temperature dependent turbidity assays and differential interference contrast (DIC) microscopy. Moreover, we unravel the molecular mechanism underlying the QW LLPS and construct the temperature-dependent density profiles (i.e., the phase diagram). To the best of our knowledge, this is the first study demonstrating the LLPS behavior of dipeptides (the minimalistic peptide) which is intractable by existing theoretical and computational methods for studying protein LLPS.

## Results

### The aggregation capability of each of the 400 dipeptides

For each of the 400 dipeptides, we have performed 1~2 individual micro-second simulations starting from 720 dipeptide molecules randomly dispersed in aqueous solution. Reaching equilibrium is essential to characterizing the phase behavior of peptides, so we assess the convergence of each simulation by tracking the time evolution of solution accessible surface area (SASA) (Fig. S4), and the length of each simulation varies from 2.5~10.0 μs (the number of simulations for each dipeptide as well as the time length of each simulation are given in Table S1-3).

An important feature of LLPS is the separation of a solute-rich dense phase from the homogenous solution, thus the first step in deciphering the LLPS propensity of a dipeptide is to identify its aggregation capability. Tuttle and Ulijn have performed short time-scale MD simulations on all 400 dipeptides, and screened out dipeptides with high self-assembly tendencies by defining an “aggregation propensity” (AP) parameter (41), which is calculated by the ratio of the solvent-accessible surface area (SASA) of all dipeptide molecules in the initial state to their SASA in the final configuration. Those high-AP peptides usually form solid-like aggregates (25, 27, 42), suggesting their low ability to undergo LLPS. As shown in Fig. S5a, the AP parameter is highly degenerate for relatively low aggregation capability peptides. We therefore introduce a new parameter: the clustering degree (defined by the number of dipeptide molecules in the largest cluster divided by total number of dipeptide molecules in the system), and characterize the aggregation capability of each dipeptide by two parameters (clustering degree and AP). For clarity, the AP parameter (41,43) is hereinafter referred as the collapse degree. Figure 1a shows the clustering degree of the 400 dipeptides as a function of their collapse degree. A gap exists around the collapse degree value of 3.0, dividing 400 dipeptides into two groups. The first group contains 54 dipeptides which present large values of collapse degree (3.0~7.0) and clustering degree (>0.9), suggesting that those dipeptides have high aggregation capabilities (see snapshots of YF/WW/YY). Dipeptides in the first group include: (1) dipeptides composed by aromatic residues F/W/Y, (2) dipeptides containing an F/W residue and a hydrophobic residue (H/L/I/V/M/P/C) or S/T residue (Fig. 1b). These results reaffirm that aromatic residues are crucial for the aggregation of dipeptides, and hydrophobic residues enhance the aggregation.

**Fig. 1.**
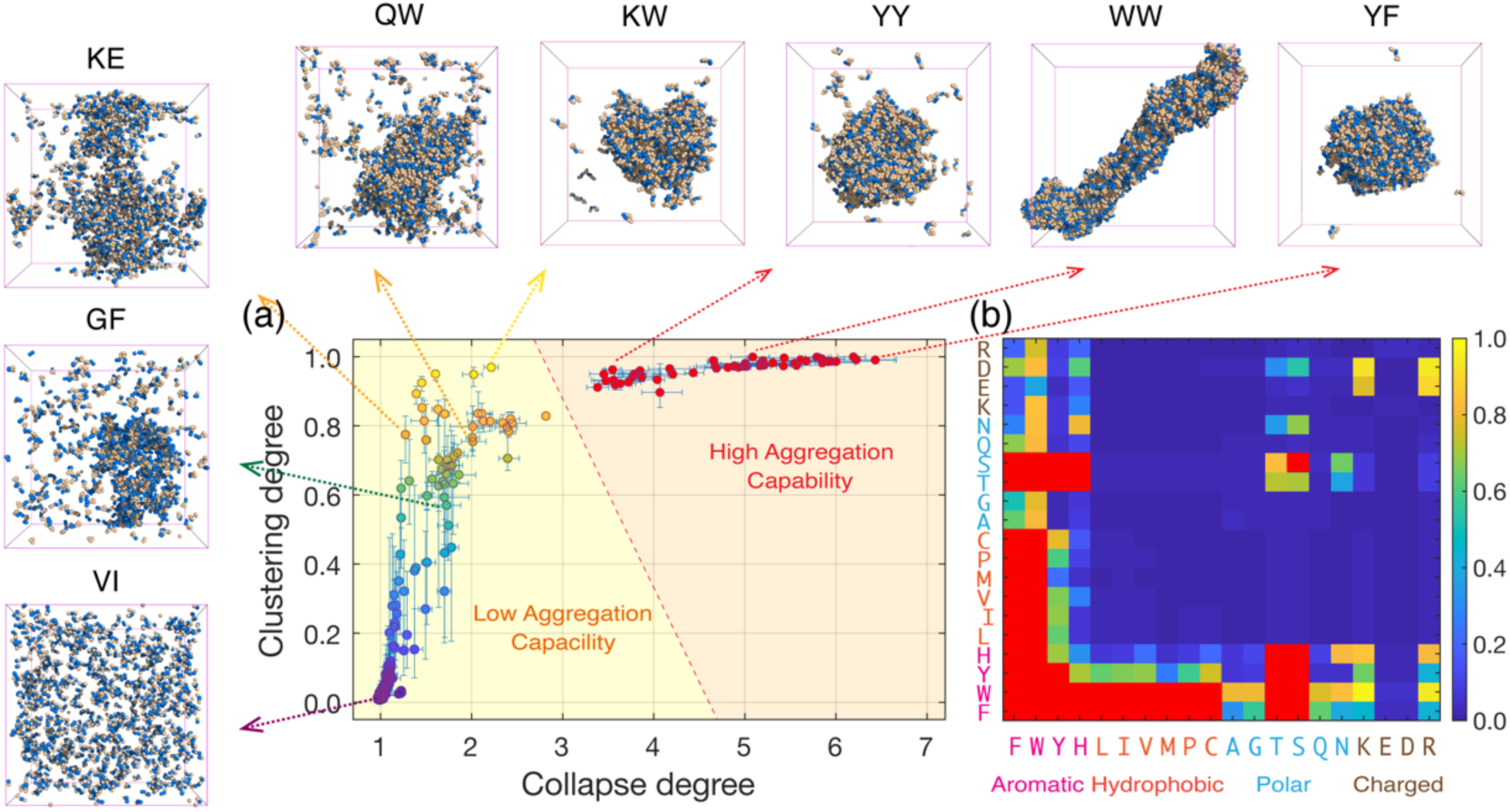
Sequence-dependent aggregation capabilities of all 400 dipeptides. (a) The clustering degree value of all dipeptides as a function of their collapse degree value. Final snapshots of 8 representative dipeptides are given. (b) The aggregation capability map of all dipeptides. Dipeptides with high aggregation capabilities are colored red, while those with relatively low aggregation capabilities are colored from blue to yellow in the order of increasing clustering degree values. The horizontal and vertical axes correspond respectively to the N- and C-terminal residue.

Dipeptides in the second group have smaller values of collapse degree (1.0~3.0), suggesting their relatively lower aggregation capabilities. However, they present a broad and continuous distribution of clustering degree. Dipeptides with high clustering degree values can aggregate into a large but loosely-packed cluster (see snapshots of WA/QW/KE/GF), while those with low clustering degree values cannot aggregate at all (see snapshots of VI). Therefore, this parameter can be used to further estimate the aggregation ability of dipeptides in the second group (they are colored from blue to yellow in the order of increasing clustering degree in Fig. 1b). We note that there is no gap in the clustering degree value distribution, so further classification of dipeptides in the second group requires alternative measurements.

### The fluidity of the dipeptide aggregates

Aggregation capability is an effective measurement of whether a dipeptide forms a peptide-rich dense phase, but additional parameters are needed to understand the dynamical properties of the aggregates and to distinguish between LLPS and LSPS. Based on physical principles on state of matters, liquids can be distinguished from solids by their fluidity: molecules in solids keep their specific neighborhood for a long time, and rarely exchange with their surroundings, while molecules in liquids can easily rearrange (8, 44). Thus, the liquid-like dipeptide aggregates can be distinguished from solid-like aggregates by the rapid exchange of contents between dense and dilute phase, and the rapid exchange of inter-molecular interactions within dense phase (44) (Fig. 2a). These two properties of liquid can respectively be described by the fluctuation extent of clustering degree parameter (*FE*, defined by the root mean square fluctuation of the dense phase size) and the exchange rate of interaction parameter (ER, defined as the fraction of inter-molecular interactions within the dense phase that are lost per unit time).

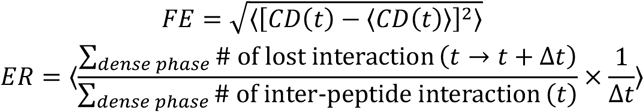

where *CD(t)* represents the clustering degree at a given time *t*, the angular bracket ⟨ ⟩ indicate averaging over conformation and over time during the last 100 ns of each simulation trajectory, and the summation is performed on all molecules within the dense phase. The *FE* and *ER* parameters reflect two independent aspect of the microscopic fluidity of the dipeptide aggregates, and they are both positively correlated with the fluidity.

**Fig. 2.**
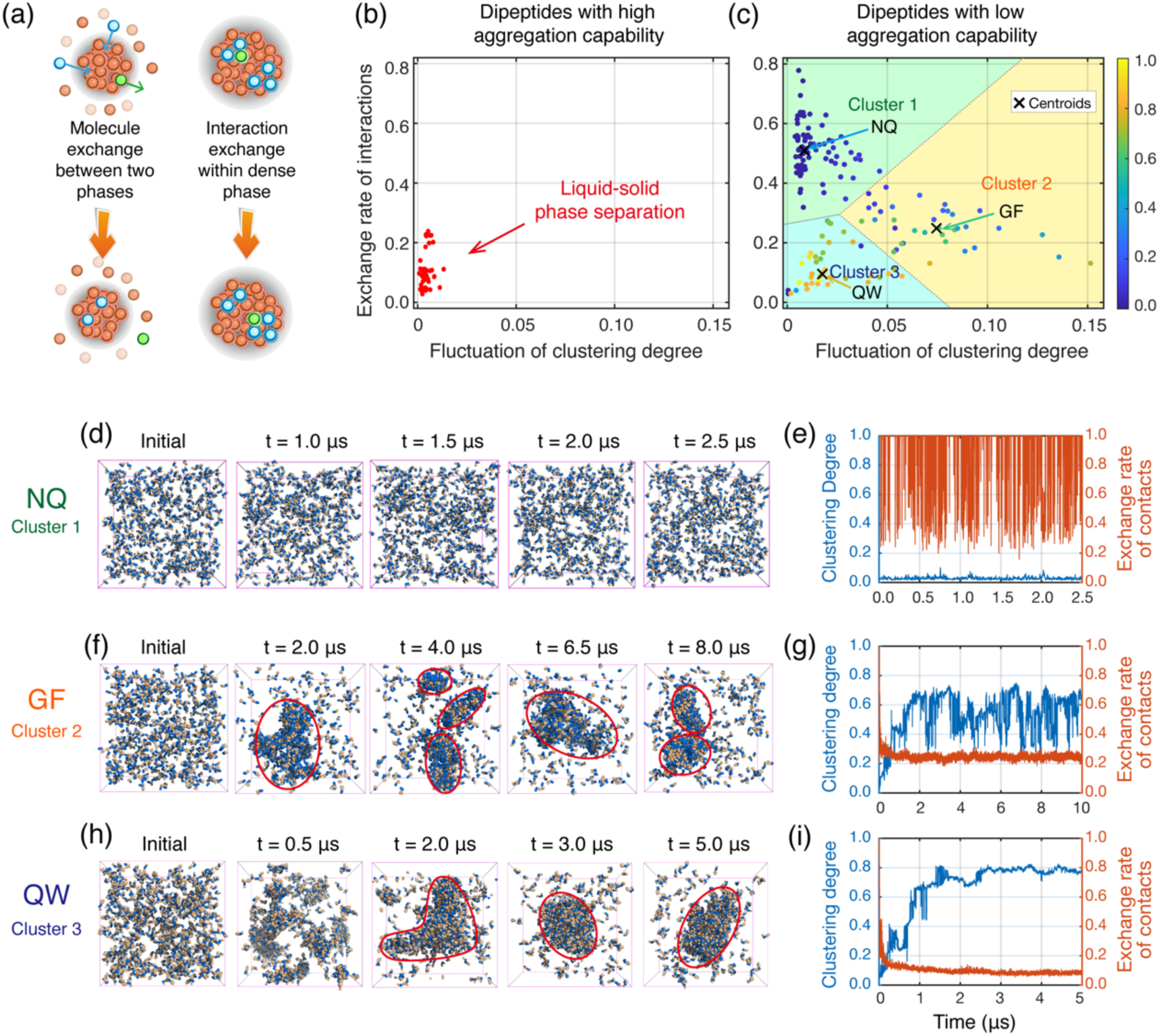
The fluidity of the dipeptide aggregates. (a) Illustration of the two parameters (the fluctuation extent of clustering degree, and the exchange rate of interactions) to characterize the fluidity of dipeptide aggregates. (b) The exchange rate of interactions values of dipeptides in group 1 (with high aggregation propensity) as a function of their fluctuation extent of clustering degree values. (c) Similar to (b) but for dipeptides in group 2 (with low aggregation propensity). The points are colored from blue to yellow in the order of increasing clustering degree values. All points are grouped into three clusters using a k-means algorithm. (d) Snapshots of NQ (closest to the centroid of cluster 1) molecules at five different time points. (e) Degree of clustering and exchange rate of contacts parameters as a function of simulation time for NQ molecules. (f-i) Similar to (d-e) but for (f-g) GF (closest to the centroid of cluster 2) molecules and (h-i) QW (closest to the centroid of cluster 3) molecules.

We calculate respectively the fluidity of dipeptides in group 1 and 2 (corresponding to dipeptides with high and low aggregation capability, respectively). Dipeptides in group 1 display small values of both *FE* and *ER*, suggesting poor fluidity and formation of solid-like aggregates (Fig. 2b and Suplementary Video 1). They are classified as dipeptides with high LSPS propensities. In contrast, dipeptides in group 2 present a broad distribution of both *FE* and *ER* values (Fig. 2c). They are further classified into three clusters using a k-means algorithm (45). NQ/GF/QW dipeptide, which is located closest to the centroid of each cluster, is taken as the representative of the corresponding cluster. NQ (centroid of cluster 1) molecules do not aggregate at all (Fig. 2d and Supplementary Video 2). Its clustering degree values maintain a small value of 0.02, and the interaction exchange rate fluctuates around a large value of 0.6 (Fig. 2e), suggesting poor phase separation ability. Interestingly, many hydrophobic dipeptides (such as VI) belong to this cluster and don’t phase separate (Supplementary Video 3). GF (centroid of cluster 2) molecules can self-assemble into a single aggregate, but the aggregate rapidly dissociates and reassociates within a microsecond timescale (Fig. 2f and Supplementary Video 4), which can be reflected by the rapid fluctuation of clustering degree (Fig. 2g). The excessive liquidity of GF aggregates suggests its inability to form liquid-like droplets. In contrast, QW (centroid of cluster 3) molecules self-associate into a single ellipsoid-like aggregate, which maintains its overall stability albeit with a certain degree of fluidity (Fig. 2h and Supplementary Video 5). The clustering degree of QW fluctuates slightly, and the *ER* value of QW maintain a much smaller value of ~0.09 compared to that of NQ (~0.6) and GF (~0.25), indicative of liquid-like QW aggregates and possibility of forming liquid-like droplets (Fig. 2i). We thus infer that QW has a high LLPS propensity.

### Experimental verifications and characterization of QW LLPS

To verify the differences in the phase behavior of dipeptides belong to different groups, we choose four dipeptides (WW, QW, GF and VI) and perform temperature dependent turbidity assay (a standard process to monitor liquid-liquid demixing (46)). We fix the peptide concentration (1.5 mM) at physiological pH=7.4 and measure the turbidity at 350 nm upon variation of the solution temperature from 5°C to 55°C (Fig. 3a-c). We detect very little turbidity for all four dipeptide solutions below 20°C, indicating the presence of dispersed monomers. With temperature increasing, the turbidity of QW gradually rises, reaches to a maximum around the temperature 37-42°C and decreases again at even higher temperature (Fig. 3a). The variation of solution turbidity can arise from LLPS or formation of other aggregates (LSPS). To identify the presence of specific phase separated state, we employ the DIC microscopy. At 5°C, the solution is clear and no evidence of liquid-demixing is identified (Fig. 3d). Upon increasing the temperature to 25°C, various sizes of liquid droplets are observed with both larger diameter ~20 μm and smaller (1 μm or below) (Fig. 3d and Supplementary video 6). The number and size of the droplets increase with the temperature rising to 37°C, in agreement with the observed high turbidity (Fig. 3d and Supplementary video 7). However, further increase of temperature (65°C) results in disappearance of the droplets after 1h but reappear on cooling down the solution to 37°C for 3 h, indicative of temperature dependent LLPS of QW (Fig. 3d). We study the dynamics of peptide molecules in the droplets during their formation and maturation over time. A time-lapse detailed inspection of the droplets shows the coalescence of two droplets to form single larger droplets through droplet fusion, confirming their liquid-like nature (Fig. 3e). The presence of peptide molecules in the liquid droplets is confirmed from the observation that the droplets converted to solid aggregates upon complete evaporation of the solvent (Fig. S6 and Supplementary video 8). On the other hand, WW shows continuous increase of turbidity with rise of temperature probably due to formation of static solid aggregates at higher temperature (Fig. 3b). Starting from clear solution at 5°C, DIC images of WW at 25°C exhibit the formation of smaller spherical assemblies with diameter below 1 μm (Fig. 3f). These assemblies gradually transform into elongated structure at higher temperature, 37°C. Further increase of temperature (65°C) results into the formation of solid-like bigger elongated aggregates, in sharp contrast to the temperature-dependent behavior of QW. The solid-nature of these aggregates is further confirmed from the fact that these structures remain undamaged and the shape persists unaltered for long time after coming out from the peptide solution (Fig. S7 and Supplementary video 9,10). Therefore, WW undergoes temperature dependent LSPS. Both the GF and VI were unable to show any kind of phase behavior depending on the solution temperature. The turbidity of the solutions slightly decreases with rise of temperature, probably due to increase of solubility at higher temperature (Fig. 3c and S8). DIC images exhibit presence of clear solution at a range of temperature from 5°C to 65°C (Fig. 3g and S8). As phase separation ability is concentration-dependent, we increase the concentration of peptide solutions by 3 times and still unable to identify any kind of structure formation in the solution. These results clearly unveil the inability of GF and VI to undergo phase separation.

**Fig. 3.**
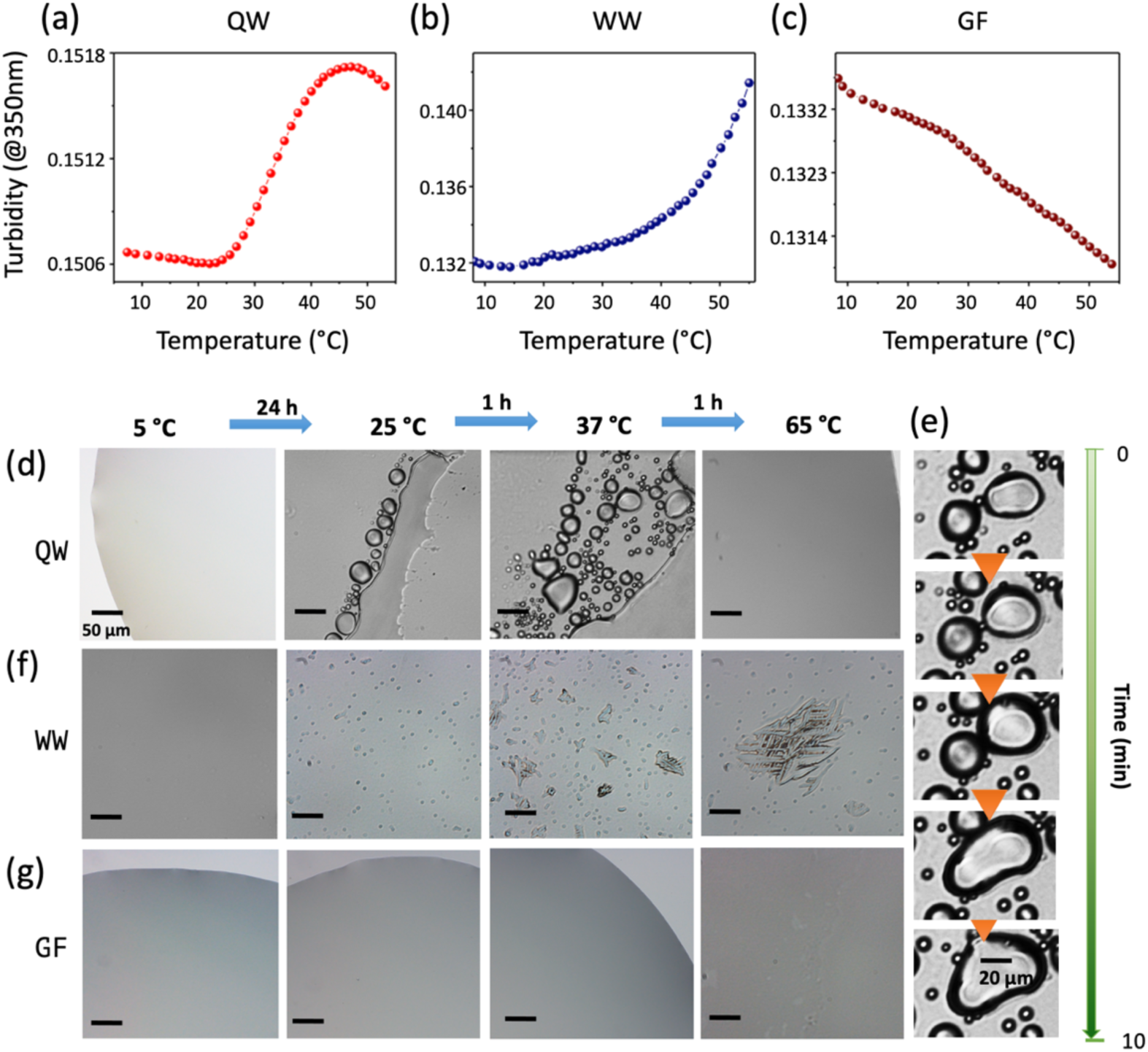
Temperature dependent phase behavior of QW, WW and GF dipeptides. (a-c) Temperature-dependent changes in turbidity (at 350 nm) of (a) QW, (b) WW, and (c) GF solutions containing 1.5 mM peptide at pH 7.4 adjusted by adding dilute HCl or NaOH (20 mM final concentration). (d-g) Temperature-dependent DIC micrographs: (d) Reversible dispersed monomer and phase-separated liquid droplets formation of QW depending on the temperature of the solution of concentration 3.01 mM at pH 7.4, (e) Time-lapse images showing fusion of two small droplets and formation of a single larger droplet over time. (f) Temperature dependent LSPS behavior of WW. (g) Images of GF solution at different temperatures indicating the inability of phase separation.

### The microscopic formation mechanism of QW liquid droplets

We perform interaction analysis to unravel the molecular mechanism underlying the QW LLPS. The QW molecule can be divided into four groups (Fig. 4a). The contact probability of each group pair in Fig. 4b shows that the Trp sidechain-sidechain pair has the highest contact probability (1.79). This interaction is characterized by the free energy landscape (FES) of Trp sidechain aromatic rings. The deep energy minimum basin located at (distance, angle) values of (0.45 nm, 10°) and the shallow basin centered at distance=0.6 nm with a widespread angle distribution (60~90°) suggest strong parallel and T-shaped aromatic stackings (Fig. 4c). The contact number between Trp/Gln mainchain and Trp sidechain (0.89/0.76) is ranked second/third. The probability density functions (PDF) of distances between Trp/Gln mainchain and Trp aromatic rings both exhibit a peak centered at ~0.65 nm (Fig. 4d-e), reflecting anion-/cation-π interactions (47). The contact probability between Gln mainchain and Trp mainchain is also high (0.47), implying hydrogen-bonding interactions. An active debate exists in literatures regarding whether proteins within liquid-like condensates form cross-β structures or remain mostly disordered (9, 15, 48–50). We examine the possible existence of β structures by calculating the FES of mainchain interactions (Fig. 4f) and observe a deep (0.7 nm, 90°) basin and a much shallower (0.5 nm, 170°) basin, indicating that QW has certain preference towards forming anti-parallel β structures, albeit with a low propensity. The solute-solvent interactions are also important for LLPS (8). Single peak is observed in the PDF curves of contact numbers between water molecules and each group in QW molecule (Fig. 4g). The ranking of peak values suggests that Gln sidechain has the highest propensity to interact with water, the Gln and Trp mainchain groups also have solvent-exposure propensity, and the aromatic Trp sidechains prefer to be buried inside the aggregate.

**Fig. 4.**
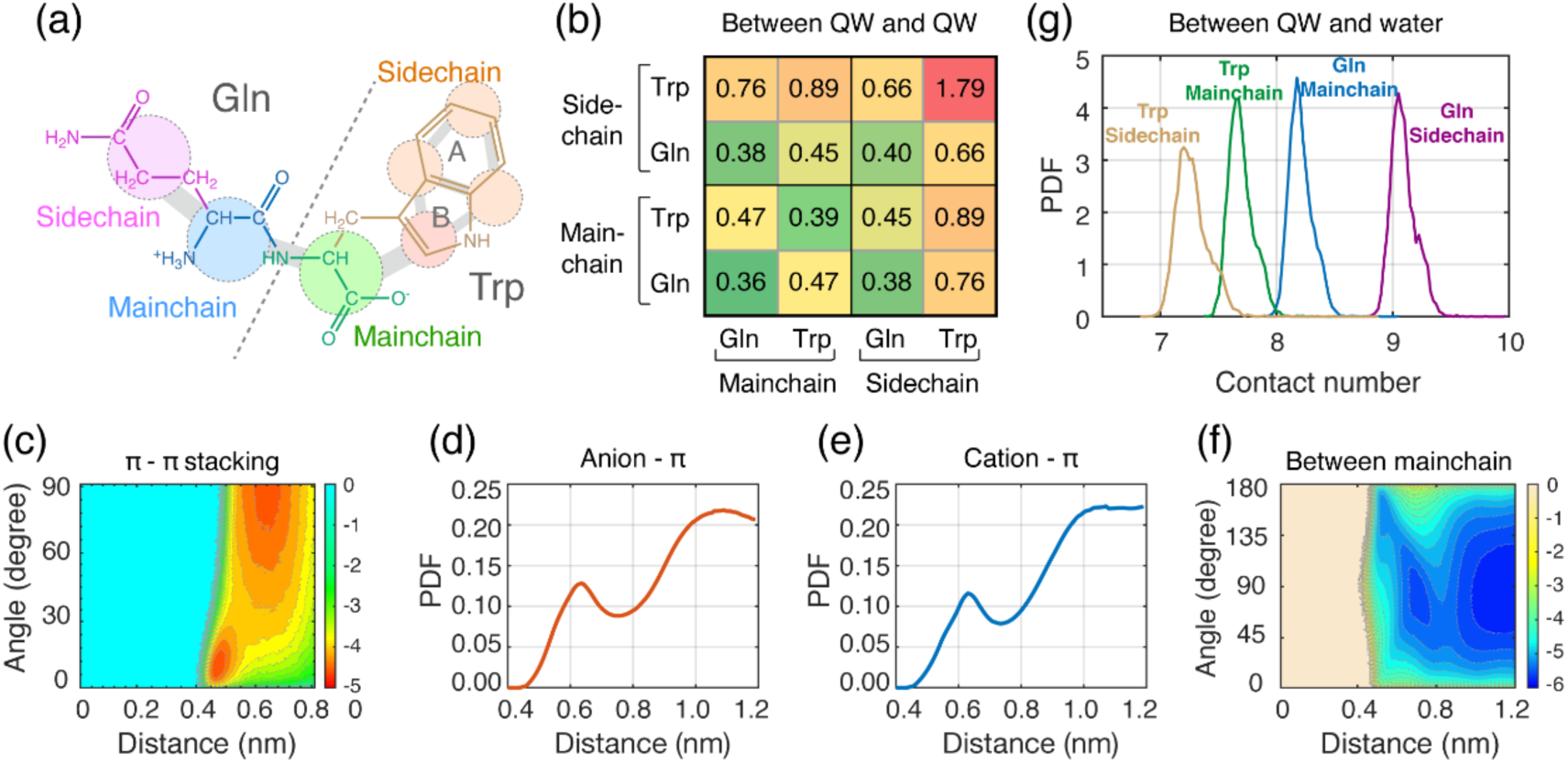
Multivalent interactions governing the LLPS of QW. (a) Mapping from all-atom model of QW molecules to the coarse-grained model. The atoms in each dashed circle are mapped to a single bead. (b) The contact probabilities or each pair of groups (mainchain/sidechain of Gln/Trp) in QW. (c) The FES as a function of centroid distance and angle between two aromatic rings of different QW molecules. (d-e) The PDF of centroid distance between Trp aromatic rings and mainchain bead of (d) Trp or (e) Gln in different QW molecules. (f) The FES as a function of centroid distance and angle between mainchains of different QW molecules. (g) The contact number between each QW group and water molecules.

Taken together, the QW LLPS is a result of the interplay among different non-covalent interactions. The inter-QW aromatic stacking interaction plays a leading role, and cooperates with the anion-/cation-π, and hydrogen-bonding interactions to drive the phase separation of QW. Meanwhile, the QW-water interaction adds to the solubility of QW, thus increase the fluidity of the aggregate. Eventually, the delicate balance between QW-QW interactions and QW-water interactions leads to the formation of liquid-like droplets.

### The temperature-dependent phase diagram of QW

Many factors including temperature, salt concentration and pH condition have been shown to affect the phase separation of diverse proteins (49, 51). Changes to the phase behavior of dipeptides in response to temperature is of key importance to gain a fundamental understanding of the thermodynamics of their phase separation. According to the phase rule, temperature and the composition of the original homogeneous mixture are sufficient for determining whether phase separation occurs, and for predicting the compositions of coexisting phases (52). However, determining LLPS phase diagram of proteins using standard sampling techniques (such as explicit-solvent MD simulation in cubic box) remains a great challenge due to the computational cost needed to reach thermodynamical equilibrium (29) and the difficulty in calculating the densities of dense/dilute phases. Existing computational approaches proposed as settlements are all based on highly-simplified (patchy particle or one-bead-per-residue) protein models in combination with slab simulation or Gibbs-ensemble simulation (33, 35) which cannot properly describe the LLPS of ultrashort peptides. Compared to proteins, the greatly reduced conformational space of dipeptide monomers enable the determination of phase equilibria for LLPS directly from micro-second MD simulations. In addition, we develop a novel and efficient protocol to determine the densities of both dense phase and dilute phase (see below for more details).

We evenly choose 26 temperature points ranging from 300 to 350 K and perform three 1.8-μs simulations at each temperature. For each simulation, we first identify the QW molecules belonging to the dense/dilute phase, calculate the total mass of peptides in the dense/dilute phase by summing up the molecular weight of the QW molecules in that phase, estimate the volume of the dense/dilute phase by a Monte Carlo sampling algorithm (53), and obtain the density of each phase by dividing its corresponding mass of peptides by volume. We then generate the coexistence curve by fitting the density profiles to the critical exponent equation(*ρ_dense_*−*ρ_dilute_* = *A*(*T*_c_−*T*)*^β^*), where β is the critical exponent which is set to 0.365 (universality class of 3D-Heisenberg model). A detailed description to the methodology is given in the Method section and supplementary information.

As shown in Fig. 5a, the densities of the dilute phase range from 20~50 mg/mL, and those of the dense phase are within 125~250 mg/mL, in agreement with the experimental determined concentrations for several proteins (the dilute/dense phase concentration for Ddx4 is within 0~50 and 160~320 mg/mL (48), and the FUS LC domain dense phase concentration is ~120 mg/mL (13)). The concentration of the dilute/dense phase increases/decreases with the rise of temperature and displays a temperature dependent phase behavior described by an upper critical solution temperature (UCST) at ~330 K. QW molecules phase separate into a well-concentrated dense phase and a dilute phase at temperatures lower than the critical point (Fig. 5b). The phase separation capability weakens as temperature increases. Around the critical temperature, the QW molecules form several small loosely-packed aggregates (Fig. 5c). Above the critical temperature, QW totally loses its phase separation ability (Fig. 5d). The UCST behavior of QW is consistent with a previous study showing that proteins with abundant polar residues generally follow an UCST phase separation(54) (the sidechain of Gln, and mainchain of Gln and Trp are all polar, and the sidechain of Trp also has a polar nitrogen atom).

**Fig. 5.**
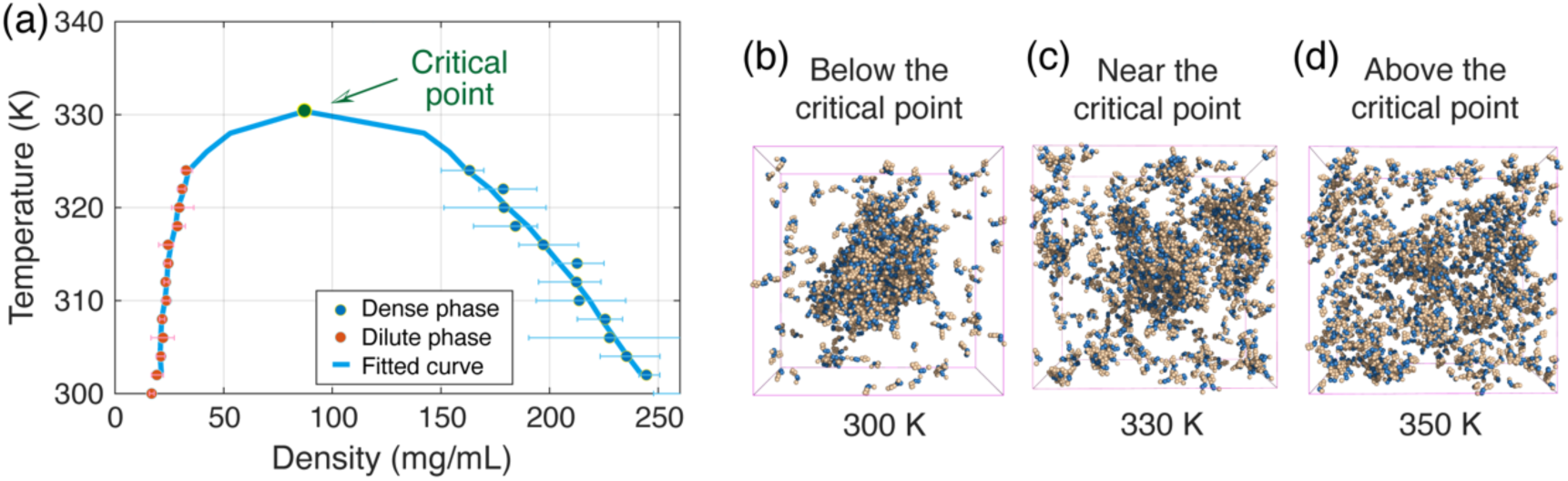
Temperature-dependent LLPS behavior of QW dipeptide. (a) The coexistence phase diagram as a function of temperature and density of the two phases. The critical point is obtained by fitting the density profiles to the critical exponent equation. (b-d) The final snapshot of a representative simulation at temperature (b) lower than, (c) near, and (d) higher than the critical temperature.

### Prediction of the LLPS propensities of all dipeptides

We predict the LLPS propensity of dipeptides (excluding the 54 LSPS dipeptides) by comparing their fluidities with QW. We define the difference between a dipeptide (A) and QW as the Euclidean distance between the (*FE*, *ER*) values of the dipeptide and those of QW:

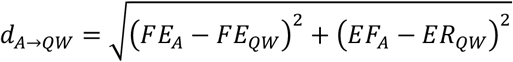

The LLPS propensity of dipeptide A 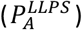 is thus defined using the equation 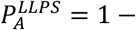 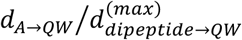, which is distributed between 0 (lowest LLPS propensity) and 1 (highest LLPS propensity). The predicted LLPS/LSPS propensities (Fig. 6a) are supported by experimental observations by us and other groups. On one hand, some dipeptides (FF, FK, FW, IF, VF, WF, WW, and WY) have been experimentally shown to form solid-like aggregates (41). They are all classified as LSPS dipeptides in our prediction. On the other hand, QW is predicted to be the most probable LLPS candidate, GF and VI are predicted to have low phase separation ability. The ability of QW and the inability of GF and VI to form liquid droplet are verified by our experiments (Fig. 3 and Fig. S8). From Fig. 6a, which shows the LLPS and LSPS propensities for all dipeptides, we propose that LLPS propensity > 0.7 indicates a good candidate for further experimental investigation. We decipher the sequence determinant of LLPS by studying the sequence characteristics of these high-LLPS-propensity dipeptides. It is apparent that aromatic residues are crucial for both LLPS and LSPS of dipeptides. Dipeptides with high LLPS propensities can be classified into three classes (Fig. 6b): 1) dipeptides composed of Y (an aromatic residue with polarity) and a hydrophobic residue L/I/V/C, 2) dipeptides composed of W (an aromatic residue with high hydrophobicity) and a polar residue A/G/Q/N (Fig. 6b), and 3) dipeptides with large dipole moment (KE/KD/RE/RD). Class 1 and 2 show that the LLPS of dipeptides requires a cooperation among aromatic stacking, hydrophobic and polar interactions. The crucial role of strong aromatic stacking interactions can be evidenced by the inability of all non-aromatic hydrophobic dipeptides to undergo LLPS, and by the lower LLPS propensities of F/H-containing dipeptides than Y/W-containing dipeptides (the π electron densities of F/H are lower than those of Y/W). The indispensability of hydrophobic and polar interactions for LLPS can be respectively shown by the inability of dipeptides composed of Y and A/G/Q/N residue to undergo LLPS, and the LSPS capability of dipeptides composed of W and L/I/V/C. Dipeptides in class 3 possess the largest dipole moment (both the mainchain and sidechain beads of the N-/C-terminal residues are positively-/negatively-charged), revealing that strong dipole-dipole interaction promotes LLPS. We notice that some dipeptides containing S and/or T residues show abnormally high LLPS/LSPS propensities compared to dipeptides containing A/G/Q/N residues. This is probably an artifact of force field, which needs to be investigated in future studies (the amber99sb/amber03ws force field parameters of S/T residues have been shown to lead to significant deviation of computational-calculated chemical shifts from experimental values (55)).

**Fig. 6.**
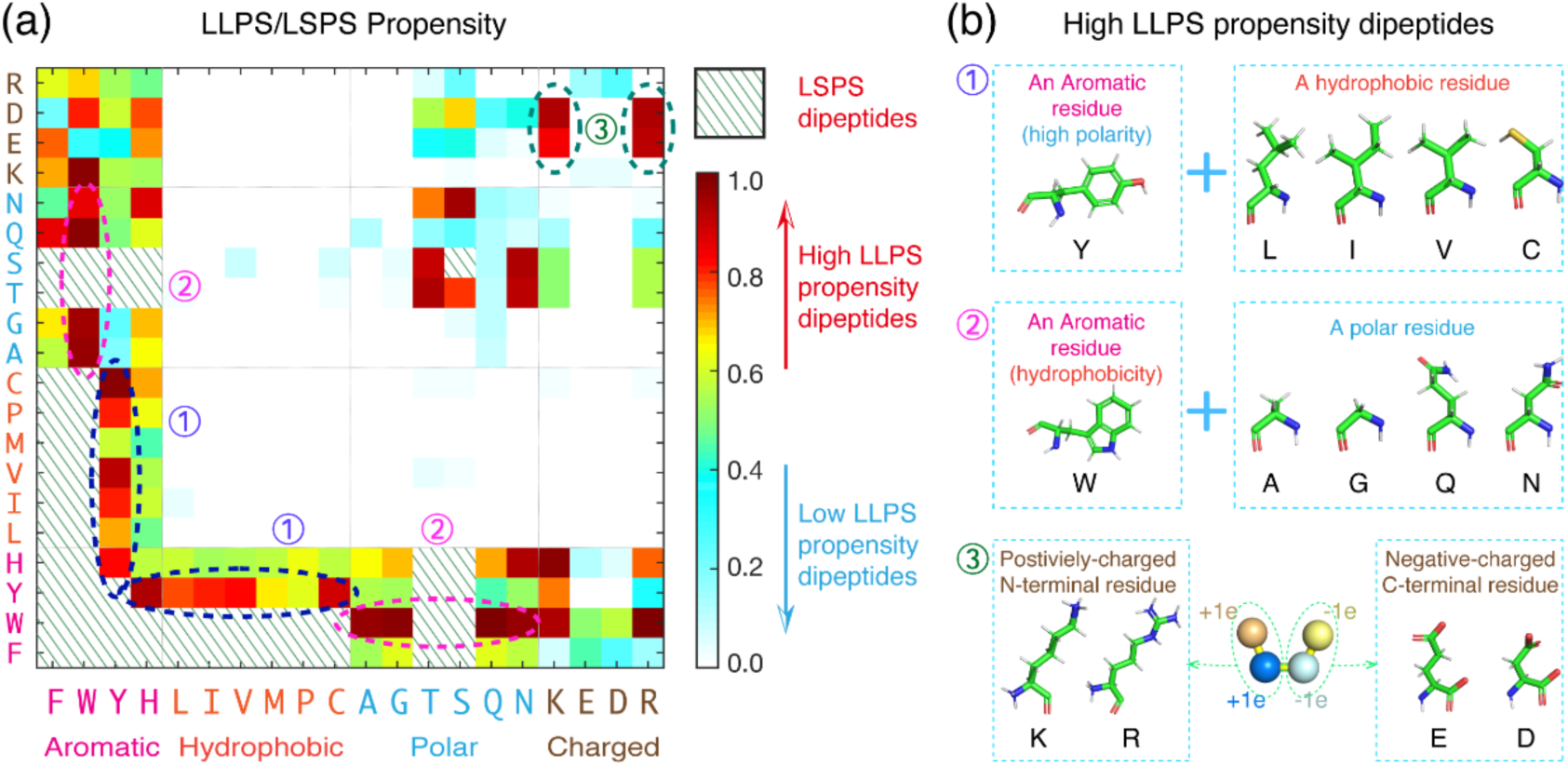
Predicted phase separation propensities of all 400 dipeptides. (a) The LSPS capability or LLPS propensity of all 400 dipeptides. The horizontal and vertical axes correspond respectively to the N- and C-terminal residue. The dipeptides with LSPS capability are represented by slash lines, while other dipeptides are colored from white to red in increasing order of LLPS propensity. The three categories of high-LLPS-propensity dipeptides are highlighted in dashed circles. (b) The illustrations to the sequence characteristics of the three categories of high-LLPS-propensity dipeptides.

Taken together, our predictions show that LLPS of dipeptides is driven by a cooperation among different types of physical interactions provided by aromatic, hydrophobic, and polar components. Compared to literature-reported proteins with LLPS capability, dipeptides have extremely short amino acid sequence lengths, but still encompass different non-covalent interaction sites, enabling them to undergo LLPS.

## Discussion

On the basis of microsecond explicit-solvent MD simulations in combination with multi-bead-per-residue coarse-grained Martini model, we comprehensively explore the LLPS abilities of all 400 dipeptide combinations of coded amino acids. A similar simulation protocol has been used by Tuttle and coworkers to predict the self-assembly propensity of dipeptides/tripeptides, albeit with a short simulation time scale. They find that aromatic dipeptides have high self-assembly propensities. Nevertheless, the LLPS ability of dipeptides and their phase behaviors remain completely elusive. We have developed a novel, efficient approach to predict the LLPS propensities of ultrashort peptides which consists of two steps: (1) determine the aggregation capability of a peptide, and (2) evaluate the fluidity of the peptide aggregates to distinguish between LLPS and LSPS. Tuttle and co-workers define the AP parameter to characterize the aggregation capability of dipeptides(41). The AP parameter, defined as the fraction of decrease in SASA, which reflects the compactness of a dipeptide aggregate, is a good quantity to screen out dipeptides with high self-assembly ability within short simulation time-scales, but is highly degenerative for low-AP dipeptides. In order to quantify aggregation capabilities of both high-AP and low-AP dipeptides, we perform micro-second-long simulations and use two parameters by introducing a new parameter, i.e., the clustering degree. On the other hand, computational characterization of the microscopic fluidity of peptides is an unexplored area. Based on physical principles on state of matters (44), we define the exchange rate of interactions and the fluctuation extent of clustering degree parameters to effectively evaluate the fluidity of the peptide aggregates.

One of the most important progresses of this work is the identification of spontaneous LLPS in dipeptide systems. We have identified 54 dipeptides with LSPS capability, and predicted the LLPS propensity of the other 346 dipeptides, among which QW possess the highest LLPS propensity. Our predictions are validated by temperature dependent turbidity assays and DIC microscopy on four representative dipeptides (WW with LSPS capacity, and QW, GF, and VI with high, medium, and low LLPS propensities, respectively). The identified QW peptide is by far the simplest LLPS system. The computational observation of dipeptide LLPS in this work is mainly attributed to the use of multi-bead-per-residue force-field and explicit solvent model, which explicitly captures key interactions (such as aromatic stacking and peptide-water interactions) driving LLPS that are missing in one-bead-per-residue models.

Another important progress of this work is the success in constructing the LLPS phase diagram using non-slab simulations. We determine the phase diagram of QW which displays a temperature dependent phase behavior described by an UCST of ~330 K. Our constructed phase diagram can better describe the temperature dependence of the densities of both dense and dilute phases mostly due to the use of explicit solvents. The protocol provided in this work can be easily applied to other short peptide systems, thus offers a way of exploring temperature dependent phase behaviors of peptides. We note that a limitation of Martini force-field is the overestimation of water freezing point (290±5 K) (56), thus this force-field cannot properly describe phase behavior at temperature around or lower than 290 K. As a result, for peptides which display a lower critical solution temperature (LCST) behavior, a more refined water model (such as polarizable Martini water) is needed. However, this will greatly increase computational costs.

In summary, our systematic investigation suggests that dipeptides, minimal but complete, possess multivalent interactions sufficient for LLPS. Three distinct categories of dipeptides are identified to have high LLPS propensities. Our results reveal that aromatic residues are crucial for a dipeptide to have LLPS capacity, and hydrophobic and polar compartments are indispensable. Our findings, together with previous reports showing the LLPS capacity of various IDPs or IDRs with varying sequence length (57–59), suggest that LLPS is a general property of peptides/proteins, independent of their length. Our protocols, which efficiently and effectively predict the LLPS propensity of dipeptides, provides a comprehensive framework for predicting and characterizing the phase behaviors of minimalistic peptides. This study set the basis for the prediction of phase behavior at the sequence level and in-depth understanding molecular mechanisms of protein phase separation.

## Materials and methods

### Molecular dynamics simulations

All dipeptide molecules are modeled using the MARTINI coarse-grained protein model (version 2.1) with positive and negative charges assigned to main-chain beads at N and C terminal, respectively. In the initial state of each MD simulation, 720 dipeptide molecules are randomly displaced in aqueous solution containing 40,000 water beads. For dipeptides with net charges, adequate number of Na+ or Cl− ions are randomly added to the solution to neutralize the system. The details of each simulation and the analysis protocols are given in the supplementary information. Graphical analysis are preformed using the PyMOL package (60).

### LLPS propensity determination

Two dipeptide molecules are considered to be in contact if their minimum distance is within 0.7 nm. The dense phase is defined as the maximized subset of all dipeptide molecules in the system where each molecule forms contact with another molecule in this subset. The aggregation propensity of each dipeptide is modeled using clustering degree and collapse degree parameters, and the fluidity of the aggregate is modeled using two parameters: the fluctuation of cluster size and the exchange rate of interactions. Definition and calculation protocols of these four parameters are given in the supplementary information. All dipeptides with low aggregation propensity are grouped into three clusters using a k-means clustering algorithm. The LLPS propensity of each of these dipeptides are defined as the additive inverse of the Euclidean distance between the (fluctuation of cluster size, exchange rate of interactions) values of this dipeptide and those of QW. The LLPS propensity is scaled between 0 (lowest LLPS propensity) and 1 (highest LLPS propensity).

### Phase diagram construction

Three independent 1.8-μs simulations are performed at each of the 26 temperature points ranging from 300 to 350 K. The dense phase is defined as the maximized subset of all dipeptide molecules in the system where each molecule forms contact with another molecule in this subset. The mass of the dense phase is calculated by summing up the molecular weight of the QW molecules in that phase. The volume of the dense phase is calculated by a refined Monte Carlo sampling algorithm (see details in supplementary information). The mass and volume of the dilute phase is calculated by subtracting the mass and volume of the total system from those of the dense phase. The critical temperature T_c_ is obtained using a similar method to the work by Mittal and coworkers (34). The densities of the dense and dilute phase (ρ_dilute_ and ρ_dense_) as functions of temperature are fitted to the equation (*ρ_dense_* − *ρ_dilute_* = *A*(*T*_c_−*T*)*^β^*), where β is the critical exponent which is set to 0.365 (universality class of 3D-Heisenberg model). The minimum fitting temperature is chosen as the physiological temperature (310 K), and the maximum fitting temperature (326 K) is chosen to maximize the coefficient of determination R^2^. The density at the critical temperature is defined as the average peptide density of the simulation at that temperature.

### Preparation of peptide solutions

Peptide solutions are prepared by dissolving the corresponding peptide at required concentration (1.5 mM or 3.01 mM or 9 mM) in double distilled water at pH 7.4 adjusted by drop wise addition of 0.1 M HCl or NaOH to a final concentration of 20 mM. The samples are first vortexed for 2 mins, followed by sonication until the peptides dissolved completely.

### Turbidity analysis and DIC microscopy

Turbidity analysis for the dipeptides is conducted by preparing solutions at concentrations of 1.5 mM in DD water at pH 7.4. Then, 200 mL aliquots are pipetted into a 96-well plate, sealed using a Breathe-Easy sealing membrane (Sigma Aldrich, Rehovot, Israel), and absorbance at 350 nm is measured. All measurements are performed using a Synergy HT plate reader (Biotek, Winooski, VT, USA) at temperature range 5°C-55°C. The heating or cooling is performed at a rate of 1 °C min−1. Temperature-dependent phase behavior of dipeptide samples is monitored by DIC microscopy. An aliquot of 15 μl peptide solutions at concentration 3.01 mM or 9 mM from said temperature are loaded onto glass coverslips and DIC images are acquired on a LSM 510 META (Zeiss) microscope with a x100 objective (oil immersion). Images are processed using ImageJ.

## Acknowledgements

This work has been supported by the National Natural Science Foundation of China (Grant Nos: 12074079 and 11674065), the National Key Research and Development Program of China (Grant Nos: 2016YFA0501702) and Israel Science Foundation (Grant No. 3145/19 to E.G). All simulations were performed using the GPU High Performance Cluster at Fudan University.

## Author Contributions

Y. T., Y. Y., and G.W. contributed to the conception of the study; Y. T. and Y. Y. performed the molecular dynamics simulations; Y. T., Y. Y., J. Z., and G.W. contributed to the data analysis and manuscript preparation. Z. L. and X. D. helped perform the analysis with constructive discussions. S. B. and E. G. performed the turbidity assay and DIC experiments. Y. T., S. B., G. W., and E. G. co-wrote the manuscript. All authors commented on the manuscript.

