## Supplemental information for "Dipeptides are minimalistic but sufficient for liquid-liquid phase separation"

### Table of contents in the supplementary information

### 1. Simulations on FF/QW/IF phase separation using HPS model + Slab box

Both experimental and computational studies have shown that FF, FW, and IF molecules can self-assemble into solid-like aggregates.[1–4] We perform Langevin Dynamics simulations on these three dipeptides using the computational framework (HPS+Slab) developed by Mittal and coworkers[5,6] to test the ability of this method to probe the phase separation of dipeptides. The peptides are described using the hydrophobicity scale (HPS) model, while the effect of water is implicitly described using a Coulombic term with Debye-Huckel electrostatic screening[7] with Debye screening length and dielectric constant set as κ = 1nm and D = 80.

In the initial state of each simulation, 720 dipeptide molecules are randomly inserted into the center of a 20*20*40 $nm^3$ simulation box (Fig. S1a, S2a, S3a). The initial equilibration is conducted for 100 ns in the NPT ensemble with periodic boundary conditions at 150 K, maintained by a Nosé-Hoover thermostat with a coupling constant of 1 ps, and 1 bar, maintained by a MTK barostat-thermostat. A time step of 10 fs is used for all the simulations. After the initial equilibration, the 720 dipeptide molecules have been evenly dispersed in the simulation box. Then the z-dimension of the box was extended to 400 nm (10 times larger than the initial z-dimension box size) while the x- and y-dimensions are kept unchanged (Fig. S1b, S2b, S3b). Next, a 100-ns-long simulation is conducted in the NVT ensemble where the temperature is gradually increased from 150 K to 300 K. With the increase of temperature, the dipeptide molecules gradually disperse from its initially well-aggregated phase. The FF and IF aggregates completely dissolve at ~180 K (Fig. S1c, S2c), and the FW aggregate completely dissolve at ~170 K(Fig. S3c), much lower than the room temperature (300 K). We further perform NVT simulations at 300 K for 4 µs, and no phase separation is observed (Fig. S1d, S2d, S3d).

Results of these simulations are not consistent with previous experimental and computational studies showing that FF, FW, IF can form solid-like aggregates, which may result from the over-simplification of the HPS model.

### 2. Methods

#### 2.1 Molecular dynamics simulations

All dipeptide molecules are modeled using MARTINI coarse-grained protein model (version 2.1)[8] with positive and negative charges assigned to main-chain beads at N and C terminal, respectively. For phase separation propensity determination, two individual simulations are performed for each of the following 92 dipeptides: The dipeptides comprising an aromatic residue F/W/Y/H and an A/G/Q/N/K/E/D/R residue, and the dipeptides comprising an aromatic residue H/Y and an H/L/I/V/M/P/C residue. (These dipeptides present moderate aggregation propensity, so two simulations are needed for obtaining statistically significant results). One simulation is performed for each of the other 308 dipeptides. The number of simulations for each dipeptide as well as the length of each simulation are shown in Table S1,2,3. For phase diagram determination for QW molecules, three independent 1.8-μs simulations are performed at each of the 26 evenly chosen temperatures ranging from 300 to 350 K.

All simulations are performed using Gromacs 2018.3 package[9]. In the initial state of each MD simulation, 720 dipeptide molecules are randomly displaced in aqueous solution containing 40,000 water beads, resulting in a size of simulation box of ~17*17*17 $nm^3$ and an effective concentration of 70~120 mg/mL. For dipeptides with net charge, adequate number of $Na^+$ or $Cl^-$ ions are randomly added to the solution to neutralize the system. Electrostatic interactions are treated using the reaction field method with a cut-off of 1.2 nm[10]. The solute and solvent are separately coupled to an external temperature bath using a velocity rescaling method[11] and a pressure bath using the Parrinello–Rahman method[12]. For the 492 simulations aiming at determining the phase separation propensity of 400 dipeptides, the temperature is maintained at 300 K. The pressure of all simulations is maintained at 1 bar. The vdW interactions are calculated using a real space cut-off of 1.2 nm. The neighborhood-list is updated every 10 steps with a cut-off distance of 1.2 nm using a Verlet buffer[13].

#### 2.2 Determining the phase separation propensity of each dipeptide

Two dipeptide molecules are considered to be in contact if their minimum distance is within 0.7 nm. The dense phase is defined as the maximized subset of all dipeptide molecules in the system where each molecule forms contact with another molecule in this subset. The aggregation propensity of each dipeptide is modeled using two parameters, degree of clustering and degree of collapse. The degree of clustering is calculated by dividing the number of dipeptide molecules in the dense phase by the total number of dipeptide molecules in the simulation box, while the degree of collapse is defined as the ratio of the solvent-accessible surface area (SASA) of all dipeptide molecules in the initial state to the SASA of them in the final configuration. The last 100 ns of each simulation trajectory is used to calculate the aggregation propensity. All dipeptides with degree of collapse parameter > 3 considered to have high aggregation propensity, and those with degree of collapse parameter < 3 have low aggregation propensity.

The fluidity of the aggregate is modeled using two parameters: the fluctuation of cluster size and the exchange rate of interactions. The fluctuation of cluster size is defined by the root mean square fluctuation of the degree of clustering parameter, and the exchange rate of interactions is defined as the fraction of inter-molecular interactions within the dense phase that are lost after 1 ns. The last 100 ns of each simulation trajectory is used to calculate the aggregation propensity. All dipeptides with low aggregation propensity are grouped into three clusters using a k-means clustering algorithm[14]. The LLPS propensity of each of these dipeptides are defined as the additive inverse of the Euclidean distance between the (fluctuation of cluster size, exchange rate of interactions) values of this dipeptide and those of QW. The LLPS propensity is scaled between 0 (lowest LLPS propensity) and 1 (highest LLPS propensity).

**2.3 Interaction analysis**

The QW molecule can be roughly divided into four groups: the positively/negatively charged mainchain group of Gln/Trp, the amphiphilic sidechain of Gln, and the aromatic sidechain of Trp. The contact number between each group and water is defined as the average number of water molecules that formed contact with this group of one QW molecule. The angle between two aromatic rings refers to the angle between the normal vectors of the two rings. If the angle is larger than 90°, the supplementary angle is used as the angle between the two phenyl rings. In the calculation, two aromatic rings are considered only if their centroid distance is within 0.8 nm. The mainchain of each QW molecule is represented by a vector pointing from its N terminal to the C terminal. The angle between the mainchains of two beads is defined as the angle between the two vectors representing these two mainchains. The two-dimensional free energy landscape is constructed using the relation $-RT\ln[P(\text{angle, centroid distance})]$, where $P$(angle, centroid distance) is the probability of a stacking pattern to have a certain value of angle and centroid distance. The distance between anion/cation and aromatic ring is defined as the distance between the negatively-/positively- charged bead (the mainchain bead of Trp/Gln) and the centroid of the aromatic ring. The data in the last 1 µs of each MD trajectory are used for interaction analysis. Trajectory visualization and graphical structure analysis are performed using the PyMOL software suite[15].

**2.4 Phase diagram construction**

Many factors including temperature, salt concentration and pH have been shown to alter the phase separation of diverse proteins[16,17]. Changes to the phase behavior of dipeptides in response to temperature is of key importance to gain a fundamental understanding of the thermodynamics of their phase separation. According to the phase rule, temperature and the composition of the original homogeneous mixture are sufficient for determining whether phase separation occurs, and for predicting the compositions of coexisting phases[18]. However, computational determination of LLPS phase diagram remains a great challenge due to the computational cost needed to reach thermodynamical equilibrium, the difficulty in calculating the densities of dense/dilute phases, and the difficulty in determining the critical temperature. To construct the phase diagram, three independent 1.8-μs simulations are performed at each of the 26 temperature points ranging from 300 to 350 K, starting from system containing 720 QW molecules randomly dispersed in water solution. The dense phase is defined as the maximized subset of all dipeptide molecules in the system where each molecule forms contact with another molecule in this subset in the last 200 ns of each simulation. The mass of the dense phase is calculated by summing up the molecular weight of the QW molecules in that phase. The volume of the dense phase is calculated by a refined Monte Carlo sampling algorithm. At least 1000 points are randomly selected in the simulation box. The neighborhood searching algorithm is performed around each of these point to determine whether a dense phase QW molecule is located within half of the interaction cutoff (0.35 nm) around this point. The volume of the dense phase equals the total volume of the simulation box times the fraction of points which has dense phase QW molecule neighbor. The mass and volume of the dilute phase is calculated by subtracting the mass and volume of the total system from those of the dense phase.

The critical temperature $T_c$ is obtained using a similar method to the work by Mittal and coworkers[5]. The densities of the dense and dilute phase ($\rho_{dilute}$ and $\rho_{dense}$) as functions of temperature are fitted to the equation$\rho_{dense} - \rho_{dilute} = A(T_c - T)^{\beta}$, where β is the critical exponent which is set to 0.365 (universality class of 3D-Heisenberg model). The minimum fitting temperature is chosen as 310 K, and the maximum fitting temperature (326 K) is chosen to maximize the coefficient of determination $R^2$. The density at the critical temperature is defined as the average peptide density of the simulation at that temperature.

**2.5 Experimental Materials**

All dipeptides were synthesized by DGpeptides Co., Ltd. (Hangzhou, China). The peptides were purified to at least 95%, and their purity and identity was confirmed by HPLC and mass spectrometry respectively.

**2.6 Preparation of peptide solutions**

Peptide solutions were prepared by dissolving the corresponding peptide at required concentration (1.5 mM or 3.01 mM or 9 mM) in double distilled water at pH 7.4 adjusted by drop wise addition of 0.1 M HCl or NaOH to a final concentration of 20 mM. The samples were first vortexed for 2 mins, followed by sonication until the peptides dissolved completely.

**2.7 Turbidity analysis**

Turbidity analysis for the dipeptides was conducted by preparing solutions at concentrations of 1.5 mM in DD water at pH 7.4. Then, 200 mL aliquots were pipetted into a 96-well plate, sealed using a Breathe-Easy sealing membrane (Sigma Aldrich, Rehovot, Israel), and absorbance at 350 nm was measured. All measurements were performed using a Synergy HT plate reader (Biotek, Winooski, VT, USA) at temperature range 5°C-55°C. The heating or cooling was performed at a rate of 1 °C min−1.

**2.8 DIC microscopy**

Temperature-dependent phase behavior of dipeptide samples was monitored by DIC microscopy. An aliquot of 15 µl peptide solutions at concentration 3.01 mM or 9 mM from said temperature were loaded onto glass coverslips and DIC images were acquired on a LSM 510 META (Zeiss) microscope with a x100 objective (oil immersion). Images were processed using ImageJ.

#### 3. Supplementary figures and tables

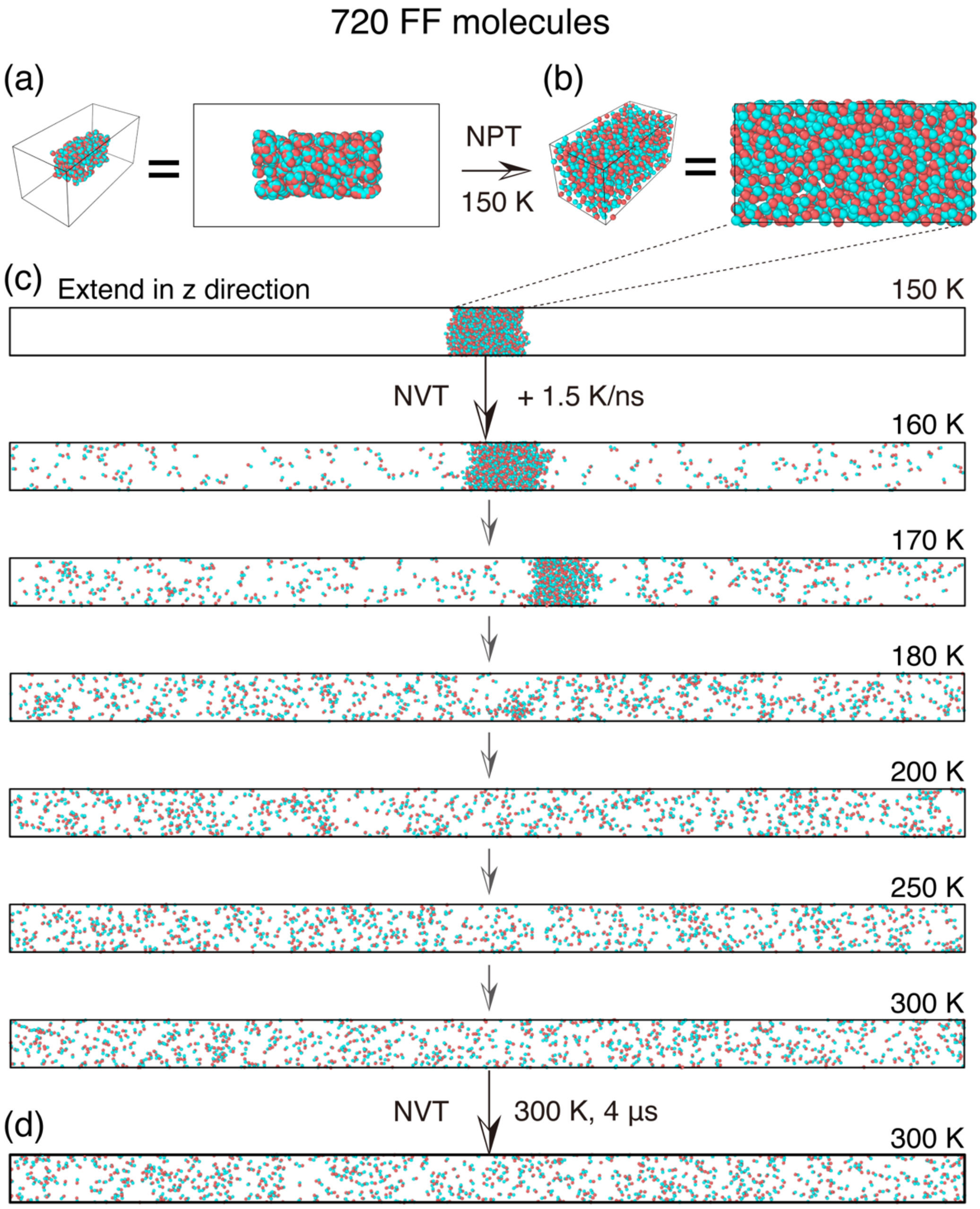

**Fig. S1. Simulations on FF molecules using HPS model and slab simulation box.**

(a) The initial state of the simulation. (b) The snapshots showing the configurations of the system after 100-ns-long NPT simulations, and the simulation box with z direction extended to 400 nm. (c) Snapshots at six different time points showing the evolution of the system with increase of temperature. (d) The snapshot of the system after 4-μs-long NVT simulation at 300 K.

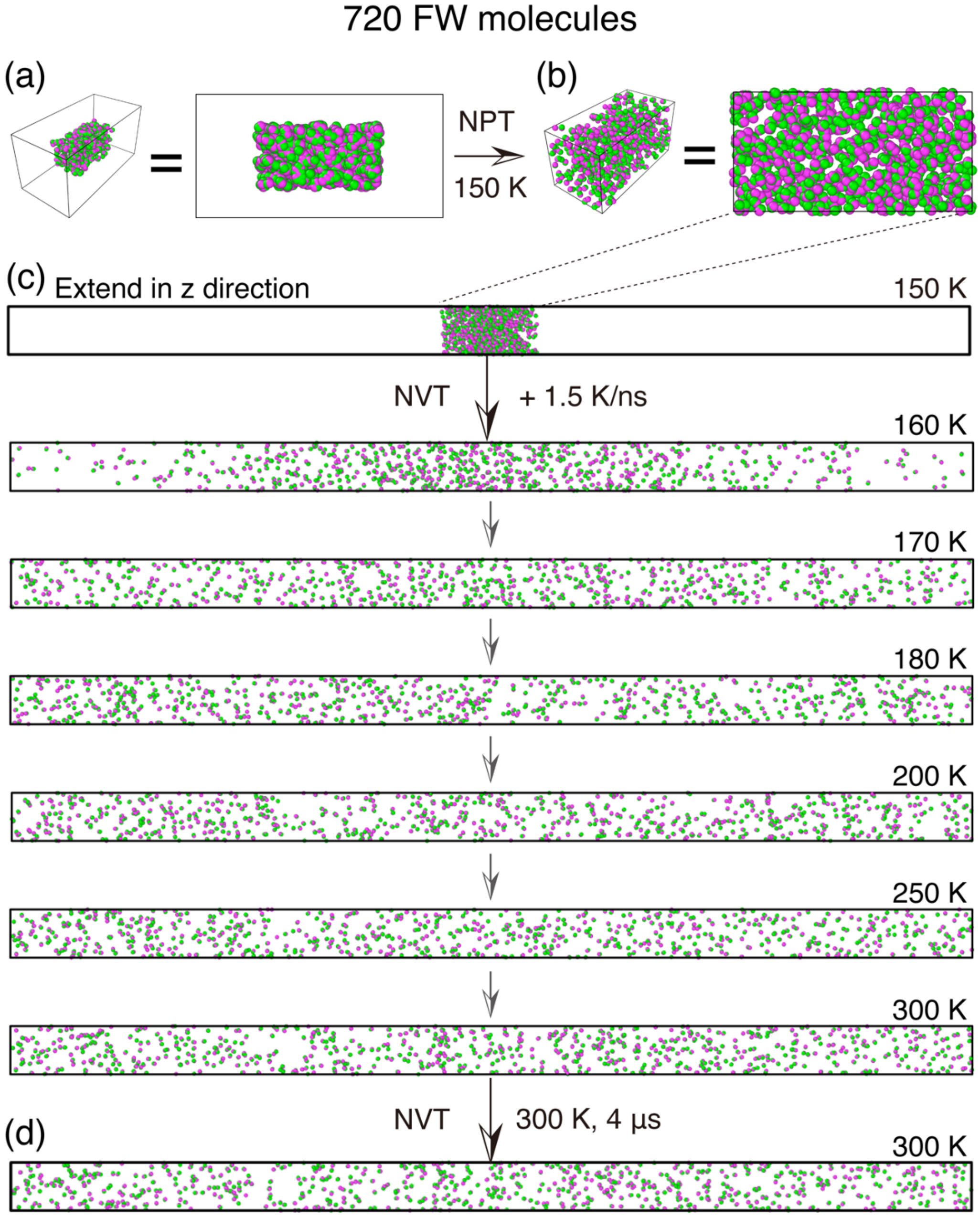

**Fig. S2. Simulations on FW molecules using HPS model and slab simulation box.**

(a) The initial state of the simulation. (b) The snapshots showing the configurations of the system after 100-ns-long NPT simulations, and the simulation box with z direction extended to 400 nm. (c) Snapshots at six different time points showing the evolution of the system with increase of temperature. (d) The snapshot of the system after 4-μs-long NVT simulation at 300 K.

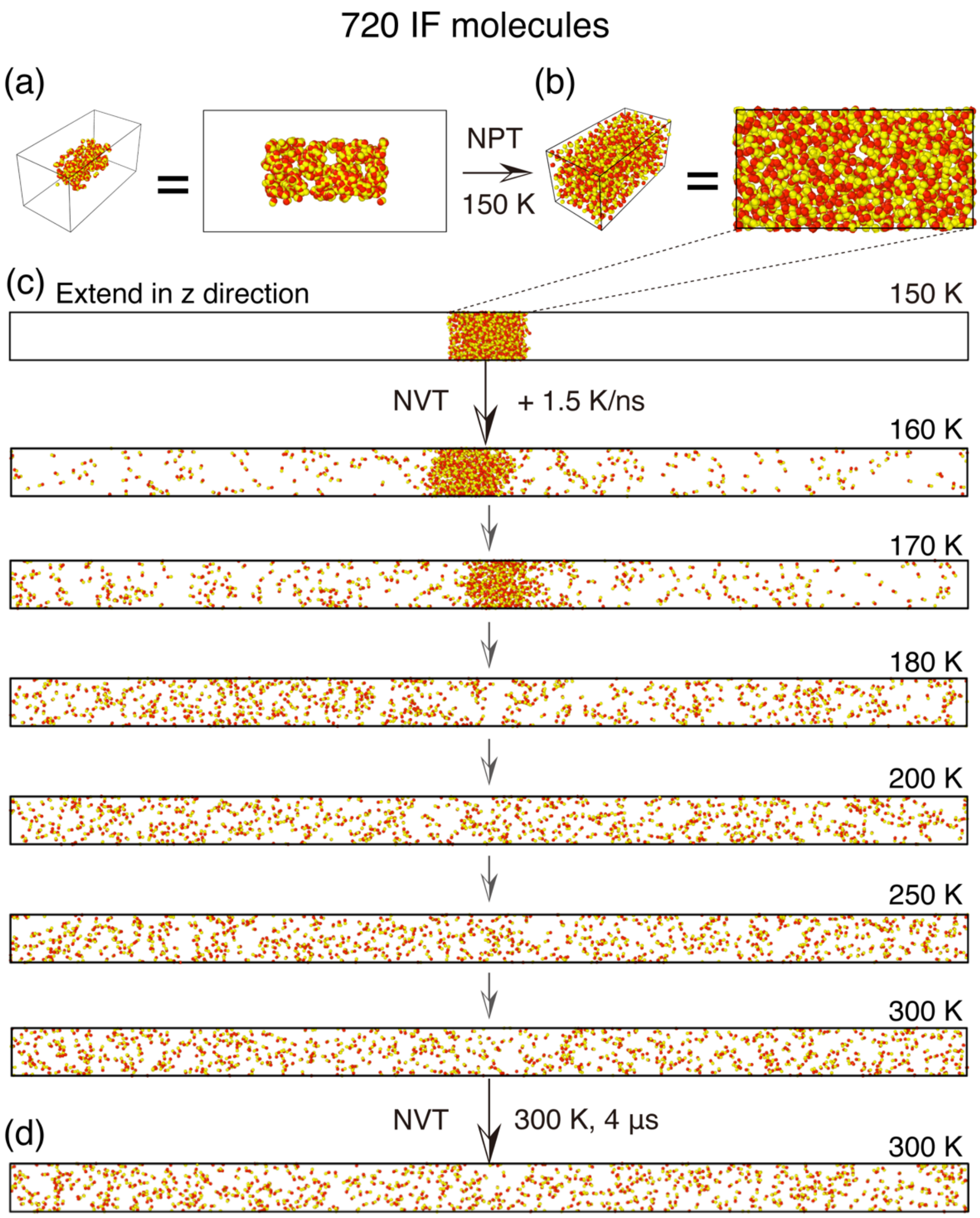

**Fig. S3. Simulations on IF molecules using HPS model and slab simulation box.**

(a) The initial state of the simulation. (b) The snapshots showing the configurations of the system after 100-ns-long NPT simulations, and the simulation box with z direction extended to 400 nm. (c) Snapshots at six different time points showing the evolution of the system with increase of temperature. (d) The snapshot of the system after 4-µs-long NVT simulation at 300 K.

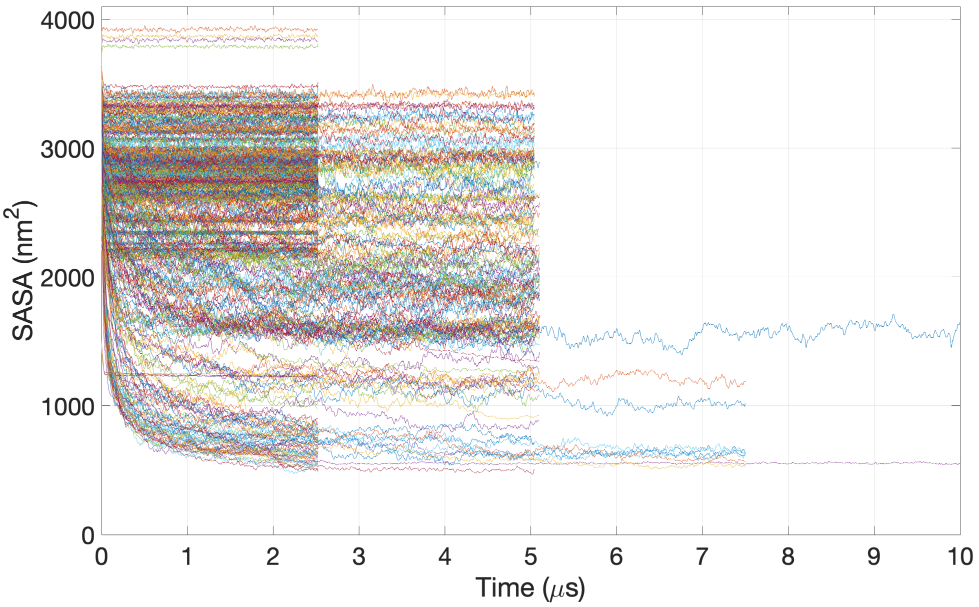

**Fig. S4. Time evolution of the solvent accessible surface area of dipeptides in all simulations.** The time length covered by the previous simulation work by Tuttle, Ulijn and their co-workers[19] is represented by an orange dashed line.

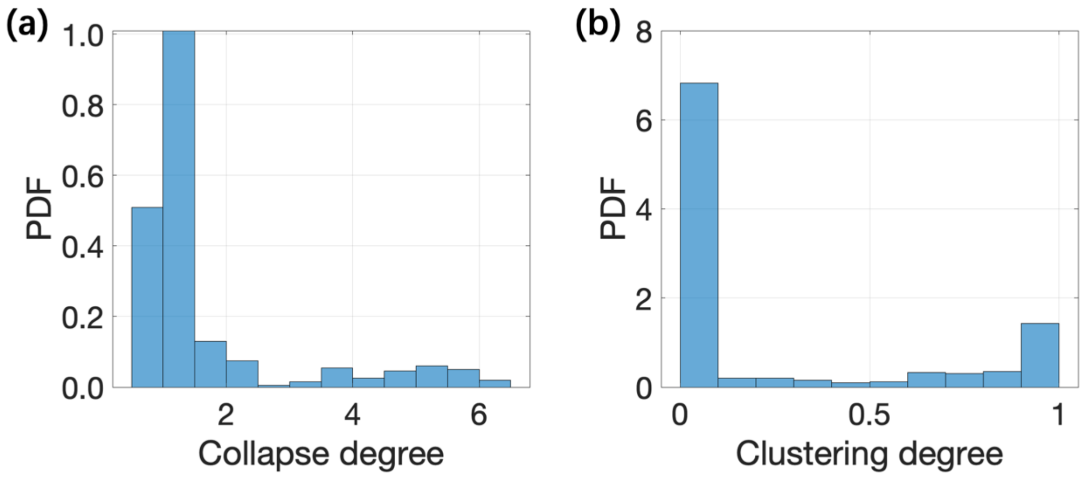

**Fig. S5. Probability density function (PDF) of the two aggregation capability parameters**
(a) PDF of the collapse degree parameter. (b) PDF of the clustering degree parameter.

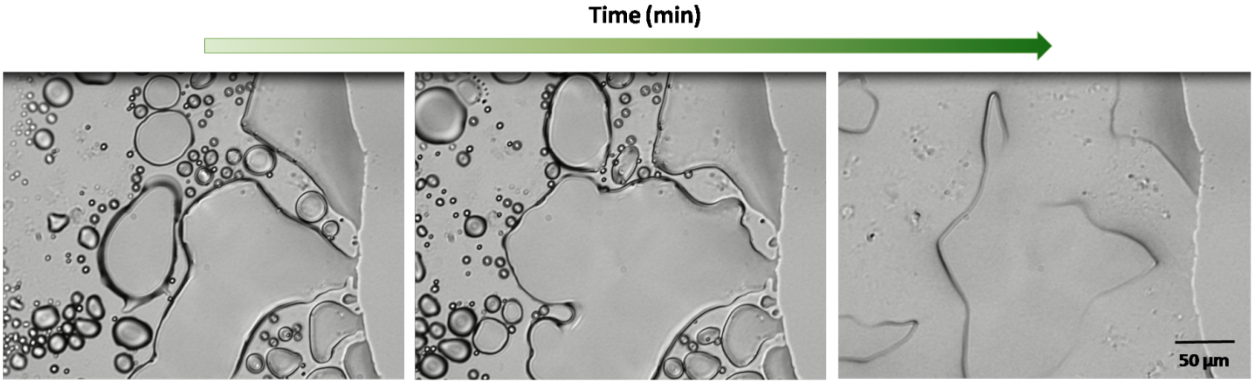

**Fig. S6. Upon complete solvent evaporation, the liquid droplets of QW converted into solid aggregates.**

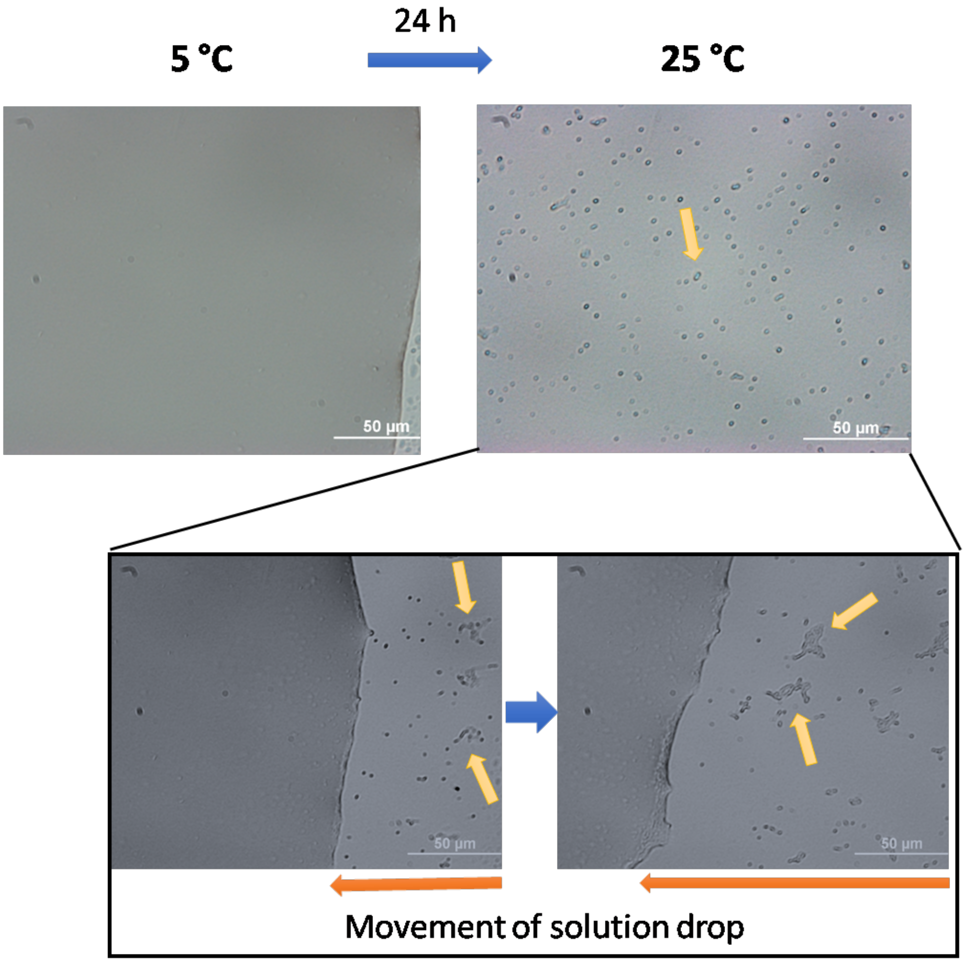

**Fig. S7. Temperature dependent LSPS of WW.**

Top right image is showing the small spherical assemblies (marked by yellow arrow) inside solution drop at 25°C. The bottom panel is showing time-lapse characteristics. As the solution drop moves (indicating by orange arrow), the spherical assemblies are coming out from solution and remain intact for long time confirming their solid-like nature.

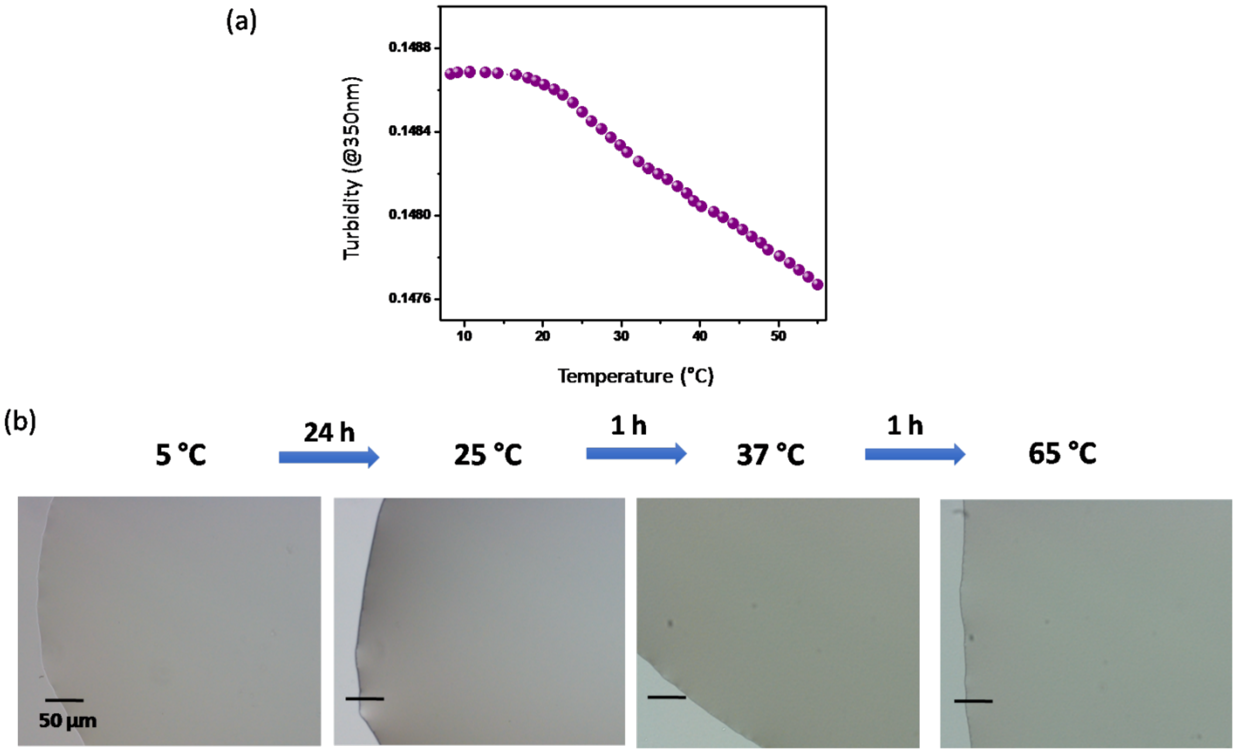

**Fig. S8. Temperature dependent behavior of VI.**

(a) Temperature dependent turbidity assay of VI solution of concentration 1.5 mM at pH 7.4. (b) DIC micrographs of 3.01 mM concentrated VI solution at different temperature emphasizing the inability of undergoing any phase separation.

**Table S1. Number of individual simulations performed for each dipeptide**

Rows: Residue at the C terminal; Columns: Residue at the N terminal

| | F | W | Y | H | L | I | V | M | P | C | A | G | T | S | Q | N | K | E | D | R |
|---|---|---|---|---|---|---|---|---|---|---|---|---|---|---|---|---|---|---|---|---|
| R | 2 | 2 | 2 | 2 | 1 | 1 | 1 | 1 | 1 | 1 | 1 | 1 | 1 | 1 | 1 | 1 | 1 | 1 | 1 | 1 |
| D | 2 | 2 | 2 | 2 | 1 | 1 | 1 | 1 | 1 | 1 | 1 | 1 | 1 | 1 | 1 | 1 | 1 | 1 | 1 | 1 |
| E | 2 | 2 | 2 | 2 | 1 | 1 | 1 | 1 | 1 | 1 | 1 | 1 | 1 | 1 | 1 | 1 | 1 | 1 | 1 | 1 |
| K | 2 | 2 | 2 | 2 | 1 | 1 | 1 | 1 | 1 | 1 | 1 | 1 | 1 | 1 | 1 | 1 | 1 | 1 | 1 | 1 |
| N | 2 | 2 | 2 | 2 | 1 | 1 | 1 | 1 | 1 | 1 | 1 | 1 | 1 | 1 | 1 | 1 | 1 | 1 | 1 | 1 |
| Q | 2 | 2 | 2 | 2 | 1 | 1 | 1 | 1 | 1 | 1 | 1 | 1 | 1 | 1 | 1 | 1 | 1 | 1 | 1 | 1 |
| S | 1 | 1 | 1 | 1 | 1 | 1 | 1 | 1 | 1 | 1 | 1 | 1 | 1 | 1 | 1 | 1 | 1 | 1 | 1 | 1 |
| T | 1 | 1 | 1 | 1 | 1 | 1 | 1 | 1 | 1 | 1 | 1 | 1 | 1 | 1 | 1 | 1 | 1 | 1 | 1 | 1 |
| G | 2 | 2 | 2 | 2 | 1 | 1 | 1 | 1 | 1 | 1 | 1 | 1 | 1 | 1 | 1 | 1 | 1 | 1 | 1 | 1 |
| A | 2 | 2 | 2 | 2 | 1 | 1 | 1 | 1 | 1 | 1 | 1 | 1 | 1 | 1 | 1 | 1 | 1 | 1 | 1 | 1 |
| C | 1 | 1 | 2 | 2 | 1 | 1 | 1 | 1 | 1 | 1 | 1 | 1 | 1 | 1 | 1 | 1 | 1 | 1 | 1 | 1 |
| P | 1 | 1 | 2 | 2 | 1 | 1 | 1 | 1 | 1 | 1 | 1 | 1 | 1 | 1 | 1 | 1 | 1 | 1 | 1 | 1 |
| M | 1 | 1 | 2 | 2 | 1 | 1 | 1 | 1 | 1 | 1 | 1 | 1 | 1 | 1 | 1 | 1 | 1 | 1 | 1 | 1 |
| V | 1 | 1 | 2 | 2 | 1 | 1 | 1 | 1 | 1 | 1 | 1 | 1 | 1 | 1 | 1 | 1 | 1 | 1 | 1 | 1 |
| I | 1 | 1 | 2 | 2 | 1 | 1 | 1 | 1 | 1 | 1 | 1 | 1 | 1 | 1 | 1 | 1 | 1 | 1 | 1 | 1 |
| L | 1 | 1 | 2 | 2 | 1 | 1 | 1 | 1 | 1 | 1 | 1 | 1 | 1 | 1 | 1 | 1 | 1 | 1 | 1 | 1 |
| H | 1 | 1 | 2 | 2 | 2 | 2 | 2 | 2 | 2 | 2 | 2 | 2 | 1 | 1 | 2 | 2 | 2 | 2 | 2 | 2 |
| Y | 1 | 1 | 1 | 2 | 2 | 2 | 2 | 2 | 2 | 2 | 2 | 2 | 1 | 1 | 2 | 2 | 2 | 2 | 2 | 2 |
| W | 1 | 1 | 1 | 1 | 1 | 1 | 1 | 1 | 1 | 1 | 2 | 2 | 1 | 1 | 2 | 2 | 2 | 2 | 2 | 2 |
| F | 1 | 1 | 1 | 1 | 1 | 1 | 1 | 1 | 1 | 1 | 2 | 2 | 1 | 1 | 2 | 2 | 2 | 2 | 2 | 2 |

**Table S2. Length of the first individual simulations performed for each dipeptide (μs)**

Rows: Residue at the C terminal; Columns: Residue at the N terminal

| | F | W | Y | H | L | I | V | M | P | C | A | G | T | S | Q | N | K | E | D | R |
|---|---|---|---|---|---|---|---|---|---|---|---|---|---|---|---|---|---|---|---|---|
| R | 2.5 | 5.1 | 2.5 | 2.5 | 2.5 | 2.5 | 2.5 | 2.5 | 2.5 | 2.5 | 2.5 | 2.5 | 2.5 | 2.5 | 2.5 | 2.5 | 2.5 | 2.5 | 2.5 | 2.5 |
| D | 5.1 | 5.1 | 2.5 | 5.1 | 2.5 | 2.5 | 2.5 | 2.5 | 2.5 | 2.5 | 2.5 | 2.5 | 2.5 | 5.1 | 2.5 | 2.5 | 5.1 | 2.5 | 2.5 | 5.1 |
| E | 2.5 | 2.5 | 2.5 | 2.5 | 2.5 | 2.5 | 2.5 | 2.5 | 2.5 | 2.5 | 2.5 | 2.5 | 2.5 | 2.5 | 2.5 | 2.5 | 5.1 | 2.5 | 2.5 | 5.1 |
| K | 2.5 | 5.1 | 2.5 | 2.5 | 2.5 | 2.5 | 2.5 | 2.5 | 2.5 | 2.5 | 2.5 | 2.5 | 2.5 | 2.5 | 2.5 | 2.5 | 2.5 | 2.5 | 2.5 | 2.5 |
| N | 2.5 | 5.1 | 2.5 | 5.1 | 2.5 | 2.5 | 2.5 | 2.5 | 2.5 | 2.5 | 2.5 | 2.5 | 2.5 | 5.1 | 2.5 | 2.5 | 2.5 | 2.5 | 2.5 | 2.5 |
| Q | 5.1 | 5.1 | 2.5 | 2.5 | 2.5 | 2.5 | 2.5 | 2.5 | 2.5 | 2.5 | 2.5 | 2.5 | 2.5 | 2.5 | 2.5 | 2.5 | 2.5 | 2.5 | 2.5 | 2.5 |
| S | 2.5 | 2.5 | 2.5 | 2.5 | 2.5 | 2.5 | 2.5 | 2.5 | 2.5 | 2.5 | 2.5 | 2.5 | 5.1 | 5 | 2.5 | 5.1 | 2.5 | 2.5 | 2.5 | 2.5 |
| T | 2.5 | 2.5 | 5.1 | 2.5 | 2.5 | 2.5 | 2.5 | 2.5 | 2.5 | 2.5 | 2.5 | 2.5 | 5.1 | 5.1 | 2.5 | 2.5 | 2.5 | 2.5 | 2.5 | 2.5 |
| G | 5.1 | 5.1 | 2.5 | 2.5 | 2.5 | 2.5 | 2.5 | 2.5 | 2.5 | 2.5 | 2.5 | 2.5 | 2.5 | 2.5 | 2.5 | 2.5 | 2.5 | 2.5 | 2.5 | 2.5 |
| A | 5.1 | 7.5 | 2.5 | 2.5 | 2.5 | 2.5 | 2.5 | 2.5 | 2.5 | 2.5 | 2.5 | 2.5 | 2.5 | 2.5 | 2.5 | 2.5 | 2.5 | 2.5 | 2.5 | 2.5 |
| C | 2.5 | 2.5 | 5.1 | 2.5 | 2.5 | 2.5 | 2.5 | 2.5 | 2.5 | 2.5 | 2.5 | 2.5 | 2.5 | 2.5 | 2.5 | 2.5 | 2.5 | 2.5 | 2.5 | 2.5 |
| P | 2.5 | 7.5 | 5.1 | 2.5 | 2.5 | 2.5 | 2.5 | 2.5 | 2.5 | 2.5 | 2.5 | 2.5 | 2.5 | 2.5 | 2.5 | 2.5 | 2.5 | 2.5 | 2.5 | 2.5 |
| M | 2.5 | 2.5 | 2.5 | 2.5 | 2.5 | 2.5 | 2.5 | 2.5 | 2.5 | 2.5 | 2.5 | 2.5 | 2.5 | 2.5 | 2.5 | 2.5 | 2.5 | 2.5 | 2.5 | 2.5 |
| V | 5 | 2.5 | 5.1 | 2.5 | 2.5 | 2.5 | 2.5 | 2.5 | 2.5 | 2.5 | 2.5 | 2.5 | 2.5 | 2.5 | 2.5 | 2.5 | 2.5 | 2.5 | 2.5 | 2.5 |
| I | 7.5 | 7.5 | 5.1 | 2.5 | 2.5 | 2.5 | 2.5 | 2.5 | 2.5 | 2.5 | 2.5 | 2.5 | 2.5 | 2.5 | 2.5 | 2.5 | 2.5 | 2.5 | 2.5 | 2.5 |
| L | 2.5 | 7.5 | 5.1 | 2.5 | 2.5 | 2.5 | 2.5 | 2.5 | 2.5 | 2.5 | 2.5 | 2.5 | 2.5 | 2.5 | 2.5 | 2.5 | 2.5 | 2.5 | 2.5 | 2.5 |
| H | 2.5 | 2.5 | 5.1 | 2.5 | 2.5 | 2.5 | 2.5 | 2.5 | 2.5 | 2.5 | 2.5 | 2.5 | 2.5 | 2.5 | 2.5 | 5.1 | 5.1 | 2.5 | 2.5 | 5.1 |
| Y | 2.5 | 2.5 | 7.5 | 5.1 | 5.1 | 5.1 | 5.1 | 2.5 | 5.1 | 5.1 | 2.5 | 2.5 | 2.5 | 2.5 | 2.5 | 2.5 | 5.1 | 2.5 | 2.5 | 5.1 |
| W | 2.5 | 2.5 | 2.5 | 2.5 | 2.5 | 2.5 | 2.5 | 2.5 | 2.5 | 2.5 | 5.1 | 5.1 | 2.5 | 5 | 5.1 | 5.1 | 5.1 | 2.5 | 2.5 | 5.1 |
| F | 10 | 2.5 | 7.5 | 5 | 2.5 | 2.5 | 2.5 | 2.5 | 2.5 | 7.5 | 5.1 | 10 | 2.5 | 2.5 | 5.1 | 2.5 | 5.1 | 2.5 | 2.5 | 5.1 |

**Table S3. Length of the second individual simulations performed for each dipeptide (μs)**

Residue at the C terminal (rows) / Residue at the N terminal (columns)

| | F | W | Y | H | L | I | V | M | P | C | A | G | T | S | Q | N | K | E | D | R |
|---|---|---|---|---|---|---|---|---|---|---|---|---|---|---|---|---|---|---|---|---|
| R | 2.5 | 5.1 | 2.5 | 2.5 | / | / | / | / | / | / | / | / | / | / | / | / | / | / | / | / |
| D | 5.1 | 5.1 | 2.5 | 5.1 | / | / | / | / | / | / | / | / | / | / | / | / | / | / | / | / |
| E | 2.5 | 2.5 | 2.5 | 2.5 | / | / | / | / | / | / | / | / | / | / | / | / | / | / | / | / |
| K | 2.5 | 5.1 | 2.5 | 2.5 | / | / | / | / | / | / | / | / | / | / | / | / | / | / | / | / |
| N | 2.5 | 5.1 | 2.5 | 5.1 | / | / | / | / | / | / | / | / | / | / | / | / | / | / | / | / |
| Q | 5.1 | 5.1 | 2.5 | 2.5 | / | / | / | / | / | / | / | / | / | / | / | / | / | / | / | / |
| S | / | / | / | / | / | / | / | / | / | / | / | / | / | / | / | / | / | / | / | / |
| T | / | / | / | / | / | / | / | / | / | / | / | / | / | / | / | / | / | / | / | / |
| G | 5.1 | 5.1 | 2.5 | 2.5 | / | / | / | / | / | / | / | / | / | / | / | / | / | / | / | / |
| A | 5.1 | 7.5 | 2.5 | 2.5 | / | / | / | / | / | / | / | / | / | / | / | / | / | / | / | / |
| C | / | / | 5.1 | 2.5 | / | / | / | / | / | / | / | / | / | / | / | / | / | / | / | / |
| P | / | / | 5.1 | 2.5 | / | / | / | / | / | / | / | / | / | / | / | / | / | / | / | / |
| M | / | / | 2.5 | 2.5 | / | / | / | / | / | / | / | / | / | / | / | / | / | / | / | / |
| V | / | / | 5.1 | 2.5 | / | / | / | / | / | / | / | / | / | / | / | / | / | / | / | / |
| I | / | / | 5.1 | 2.5 | / | / | / | / | / | / | / | / | / | / | / | / | / | / | / | / |
| L | / | / | 5.1 | 2.5 | / | / | / | / | / | / | / | / | / | / | / | / | / | / | / | / |
| H | / | / | 5.1 | 2.5 | 2.5 | 2.5 | 2.5 | 2.5 | 2.5 | 2.5 | 2.5 | 2.5 | / | / | 2.5 | 5.1 | 5.1 | 2.5 | 2.5 | 5.1 |
| Y | / | / | / | 5.1 | 5.1 | 5.1 | 5.1 | 2.5 | 5.1 | 5.1 | 2.5 | 2.5 | / | / | 2.5 | 2.5 | 5.1 | 2.5 | 2.5 | 5.1 |
| W | / | / | / | / | / | / | / | / | / | / | 5.1 | 5.1 | / | / | 5.1 | 5.1 | 5.1 | 2.5 | 2.5 | 5.1 |
| F | / | / | / | / | / | / | / | / | / | / | 5.1 | 10 | / | / | 5.1 | 2.5 | 5.1 | 2.5 | 2.5 | 5.1 |

### 4. Introduction to all supplementary videos

**Supplementary video 1**
Simulation trajectory showing the LSPS of WW dipeptides.

**Supplementary video 2**
Simulation trajectory showing that NQ doesn’t phase separate at all.

**Supplementary video 3**
Simulation trajectory showing that VI doesn’t phase separate at all.

**Supplementary video 4**
Simulation trajectory showing that the GF aggregates are too unstable to form liquid droplets.

**Supplementary video 5**
Simulation trajectory showing the LLPS of QW dipeptides.

**Supplementary video 6**
The DIC movis showing the droplet formation of QW at 25°C.

**Supplementary video 7**
The DIC movie showing the droplet formation of QW at 37°C.

**Supplementary video 8**
The DIC movie showing the drying process of QW droplets

**Supplementary video 9,10**
The DIC movies showing the LSPS of WW

**5. The snapshots of the final configuration of all simulations**

We have performed 491 molecular dynamics simulations to model the phase separation propensity of all 400 dipeptides. This file contains snapshots showing the final configurations of all 491 simulations. All snapshots are first sorted in decreasing order of hydrophobicity of the N-terminal residue, and then in that of the C-terminal residue.

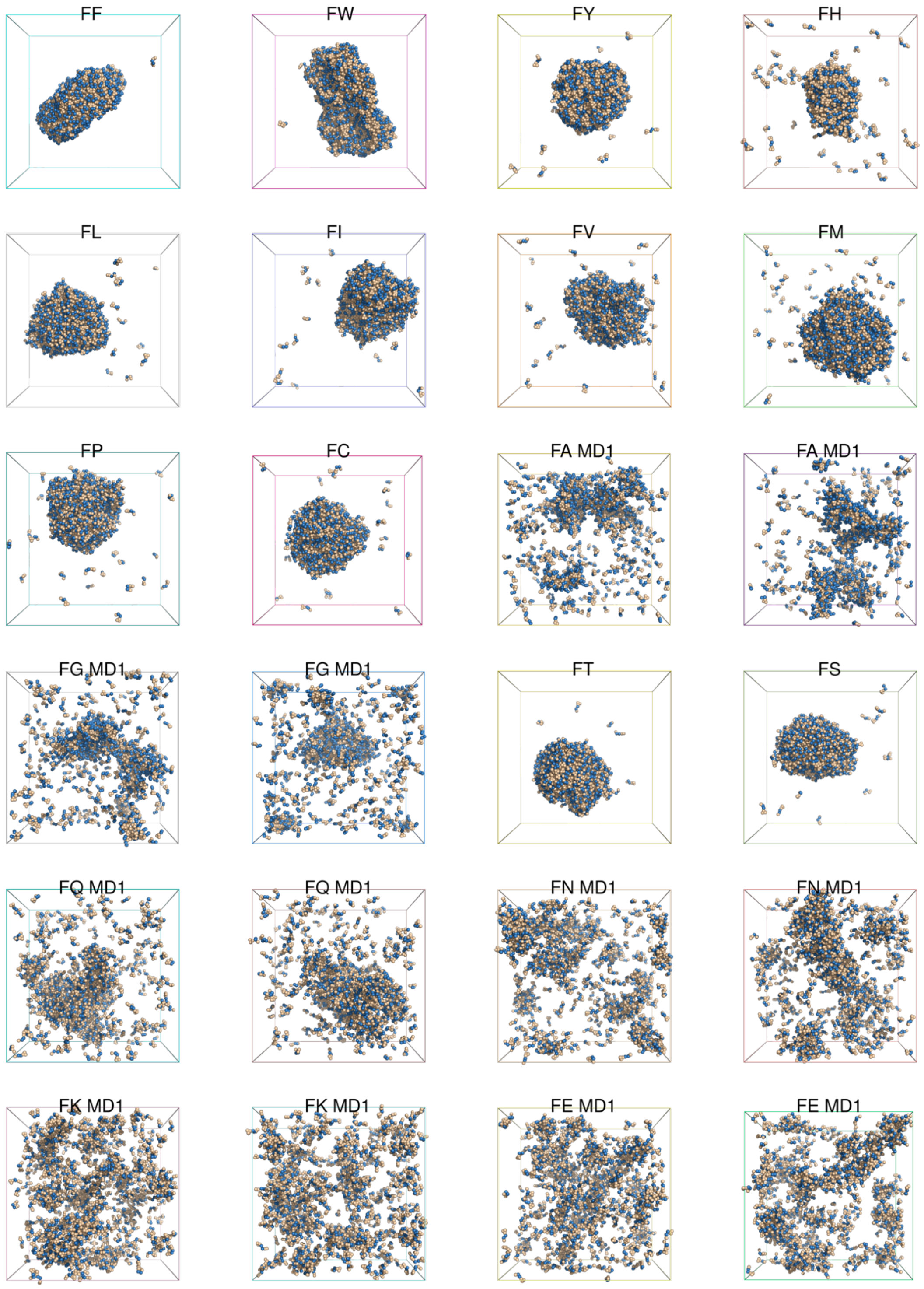

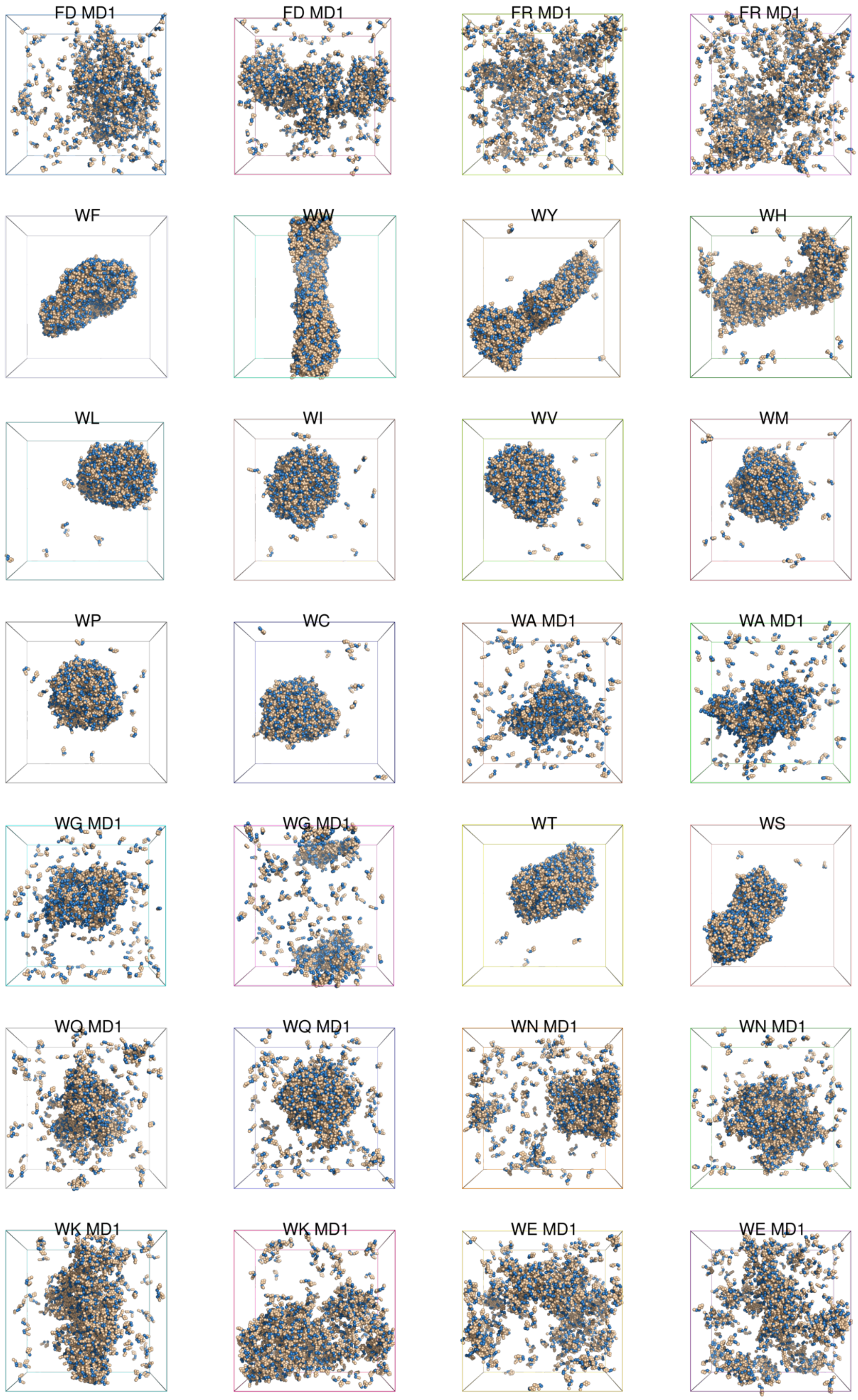
FD MD1
FD MD1
FR MD1
FR MD1
WF
WW
WY
WH
WL
WI
WV
WM
WP
WC
WA MD1
WA MD1
WG MD1
WG MD1
WT
WS
WQ MD1
WQ MD1
WN MD1
WN MD1
WK MD1
WK MD1
WE MD1
WE MD1

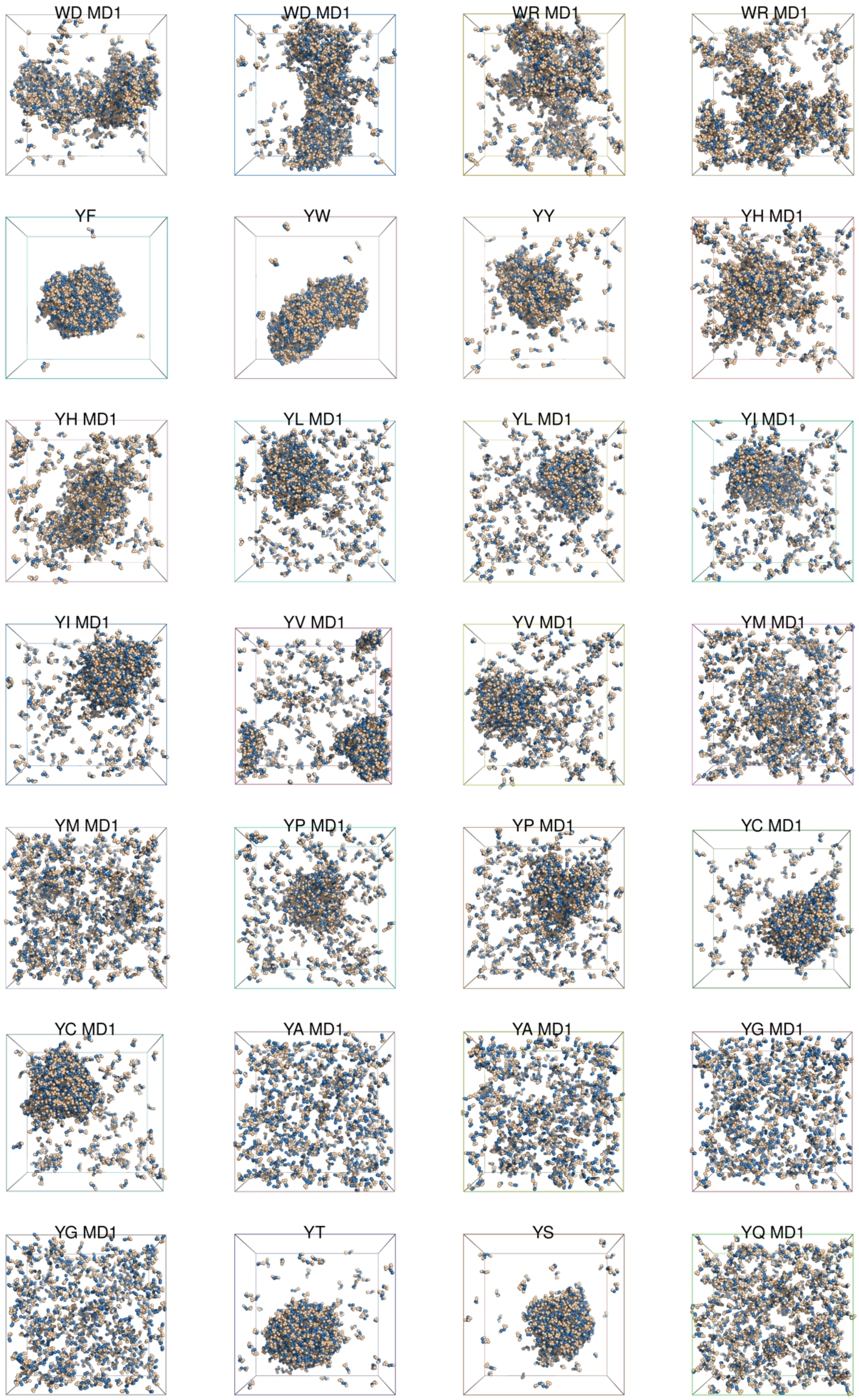
WD MD1
WD MD1
WR MD1
WR MD1
YF
YW
YY
YH MD1
YH MD1
YL MD1
YL MD1
YI MD1
YI MD1
YV MD1
YV MD1
YM MD1
YM MD1
YP MD1
YP MD1
YC MD1
YC MD1
YA MD1
YA MD1
YG MD1
YG MD1
YT
YS
YQ MD1

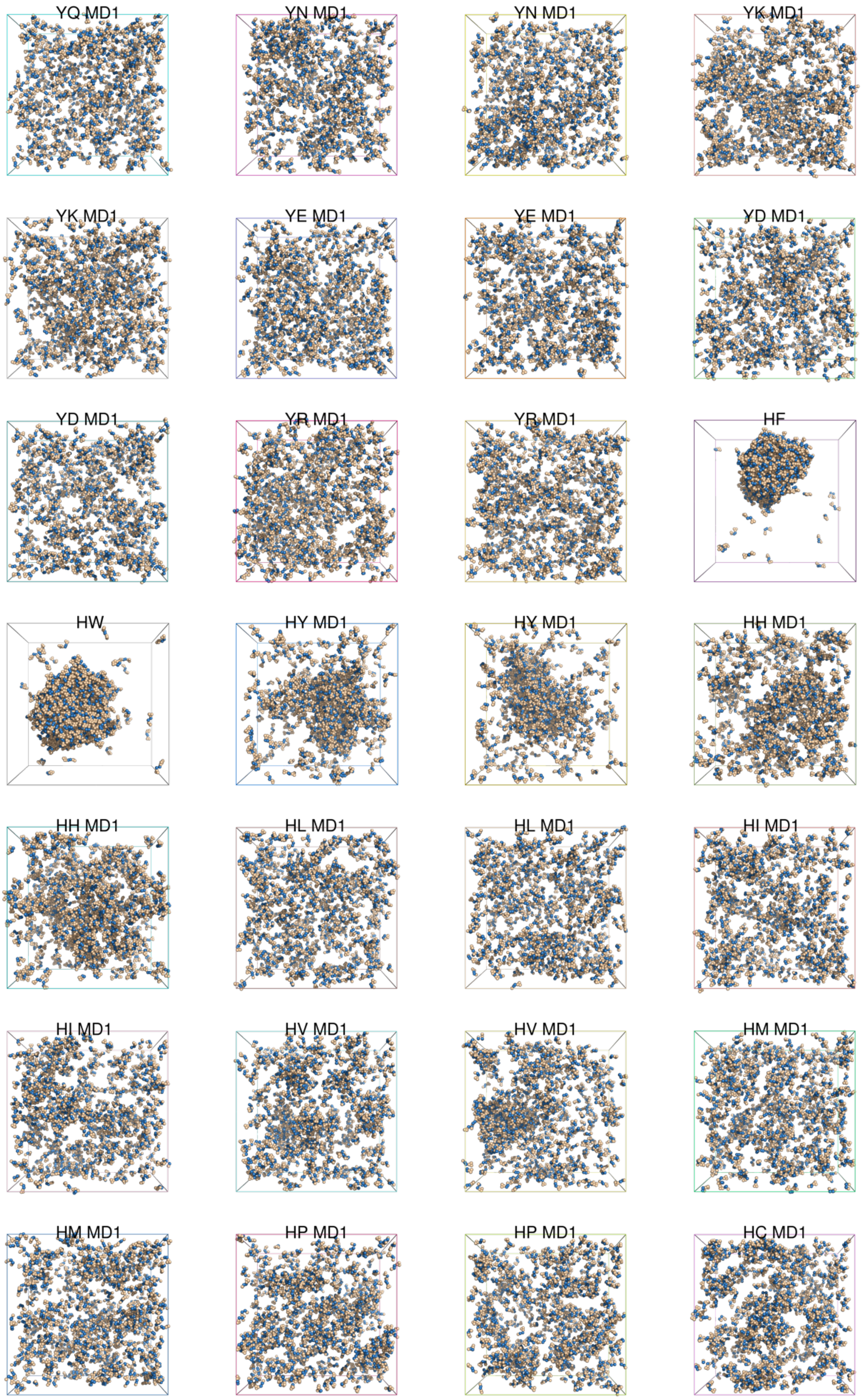
YQ MD1
YN MD1
YN MD1
YK MD1
YK MD1
YE MD1
YE MD1
YD MD1
YD MD1
YR MD1
YR MD1
HF
HW
HY MD1
HY MD1
HH MD1
HH MD1
HL MD1
HL MD1
HI MD1
HI MD1
HV MD1
HV MD1
HM MD1
HM MD1
HP MD1
HP MD1
HC MD1

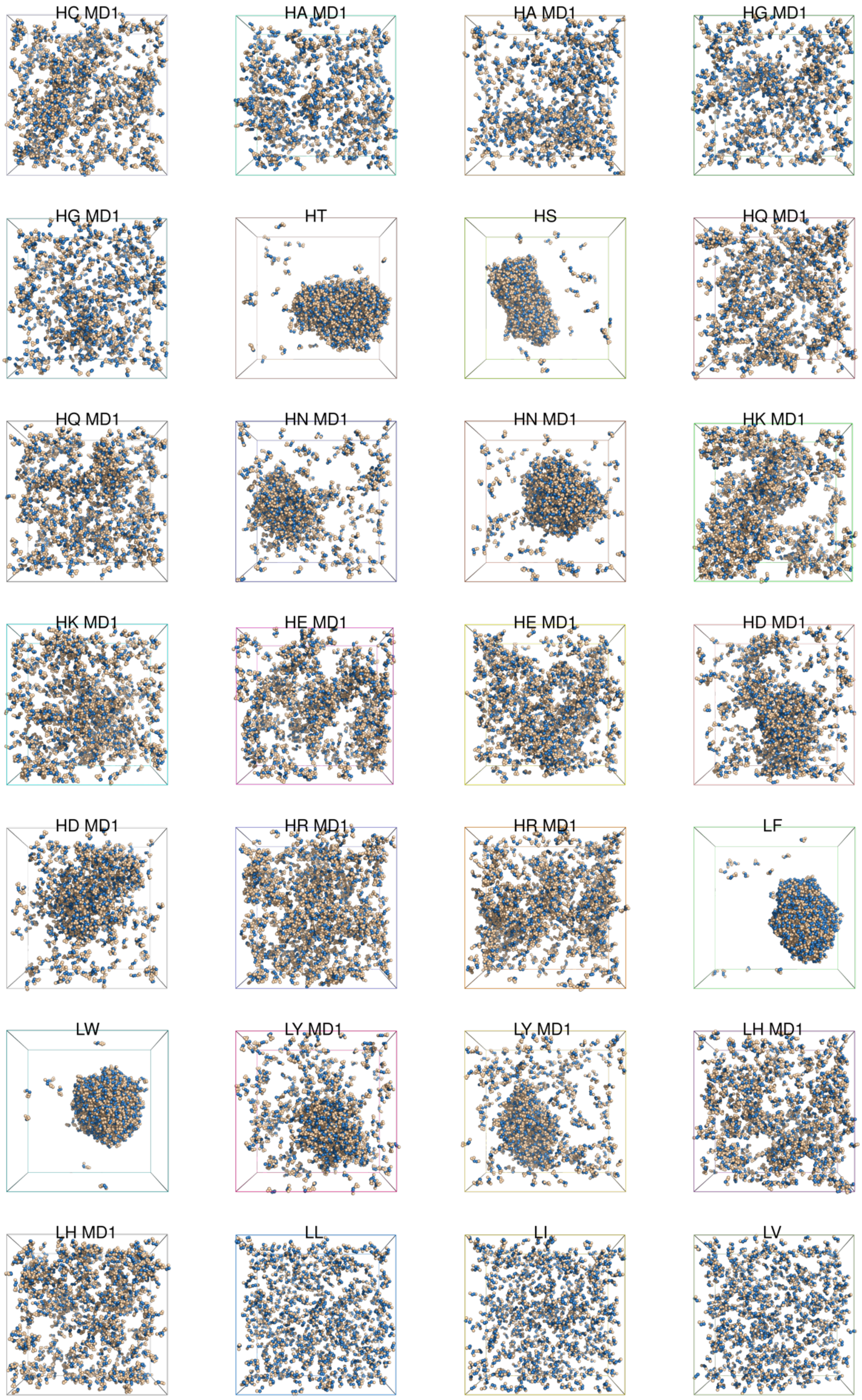
HC MD1
HA MD1
HA MD1
HG MD1
HG MD1
HT
HS
HQ MD1
HQ MD1
HN MD1
HN MD1
HK MD1
HK MD1
HE MD1
HE MD1
HD MD1
HD MD1
HR MD1
HR MD1
LF
LW
LY MD1
LY MD1
LH MD1
LH MD1
LL
LI
LV

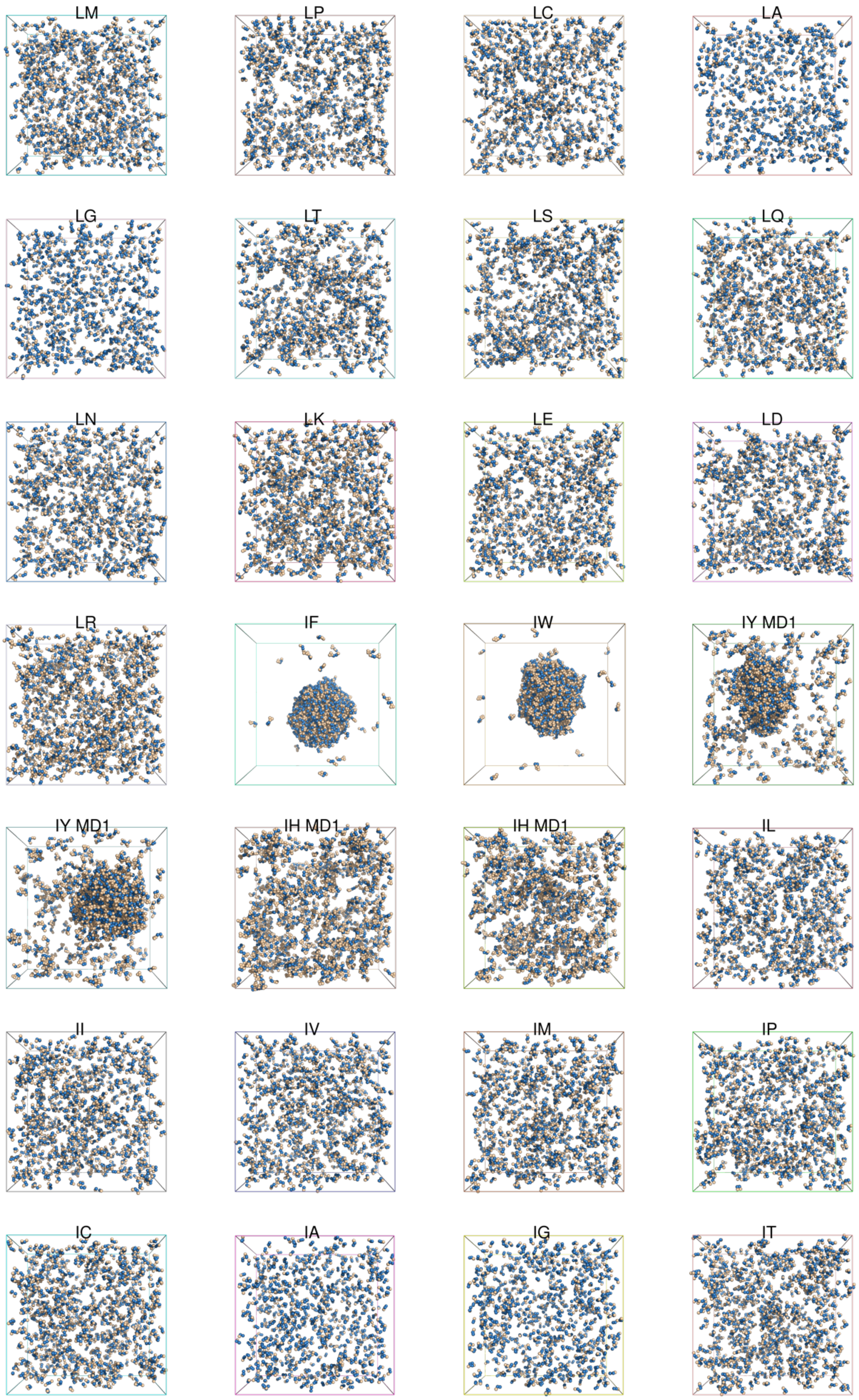
LM
LP
LC
LA
LG
LT
LS
LQ
LN
LK
LE
LD
LR
IF
IW
IY MD1
IY MD1
IH MD1
IH MD1
IL
II
IV
IM
IP
IC
IA
IG
IT

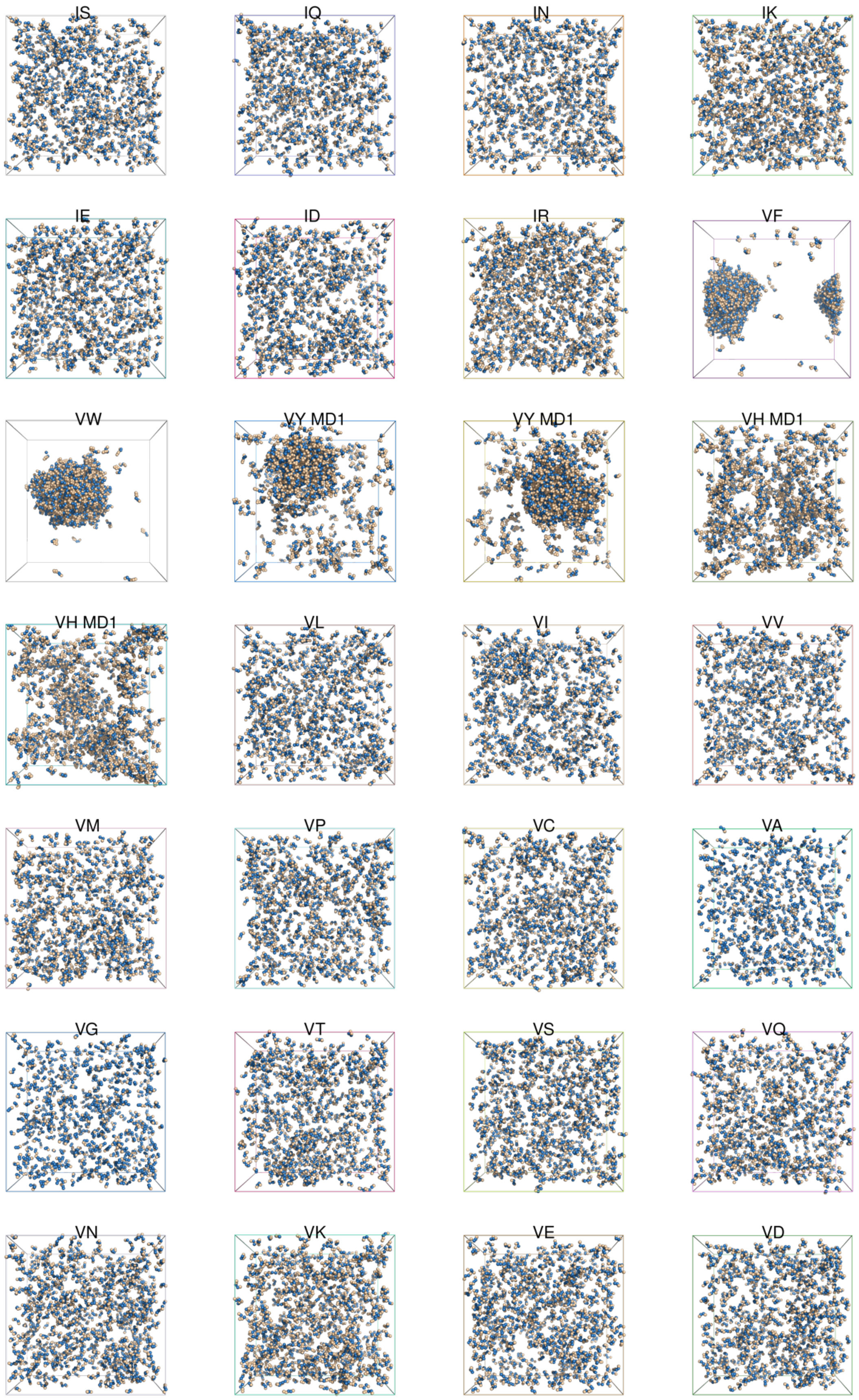
IS
IQ
IN
IK
IE
ID
IR
VF
VW
VY MD1
VY MD1
VH MD1
VH MD1
VL
VI
VV
VM
VP
VC
VA
VG
VT
VS
VQ
VN
VK
VE
VD

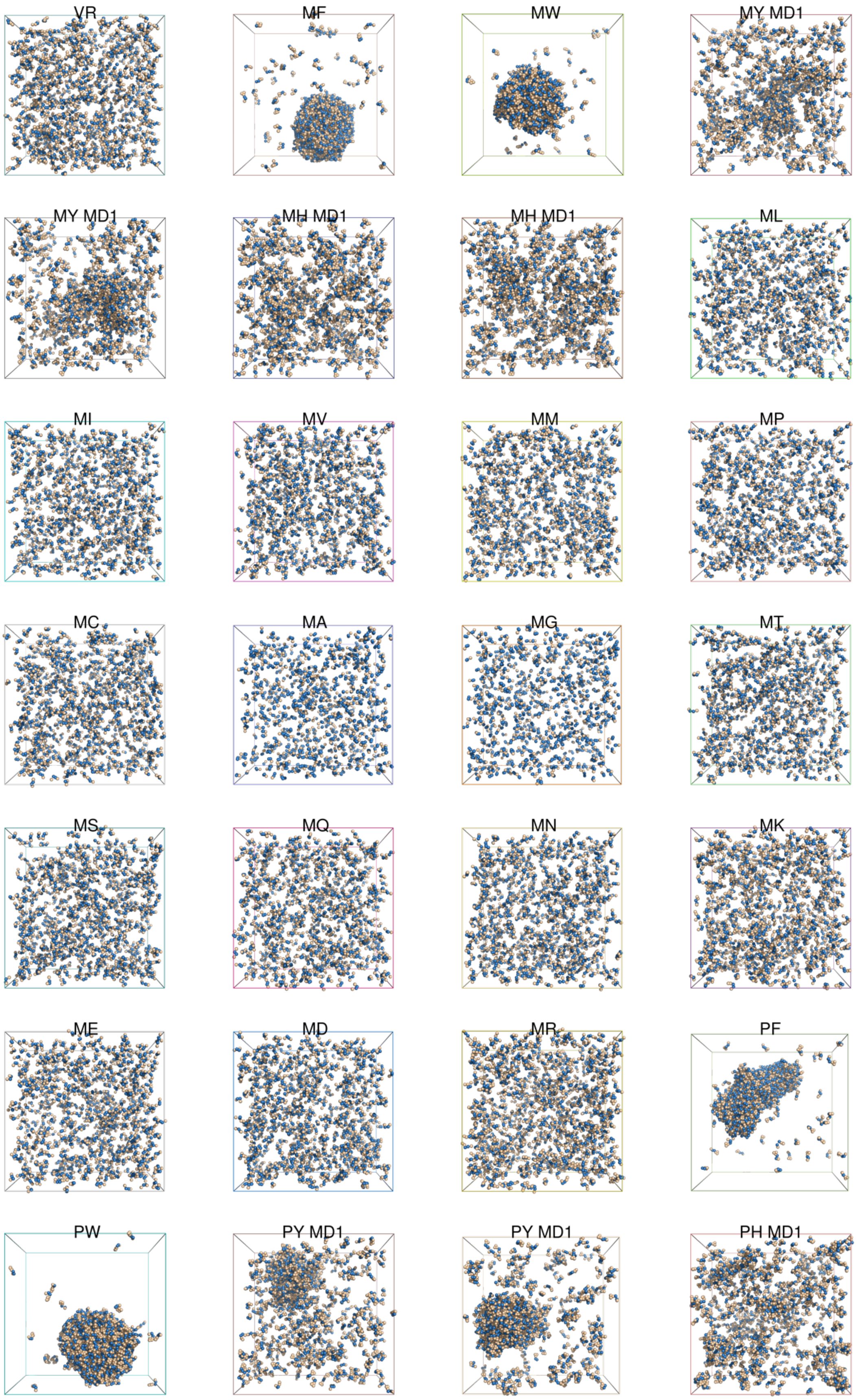
VR
MF
MW
MY MD1
MY MD1
MH MD1
MH MD1
ML
MI
MV
MM
MP
MC
MA
MG
MT
MS
MQ
MN
MK
ME
MD
MR
PF
PW
PY MD1
PY MD1
PH MD1

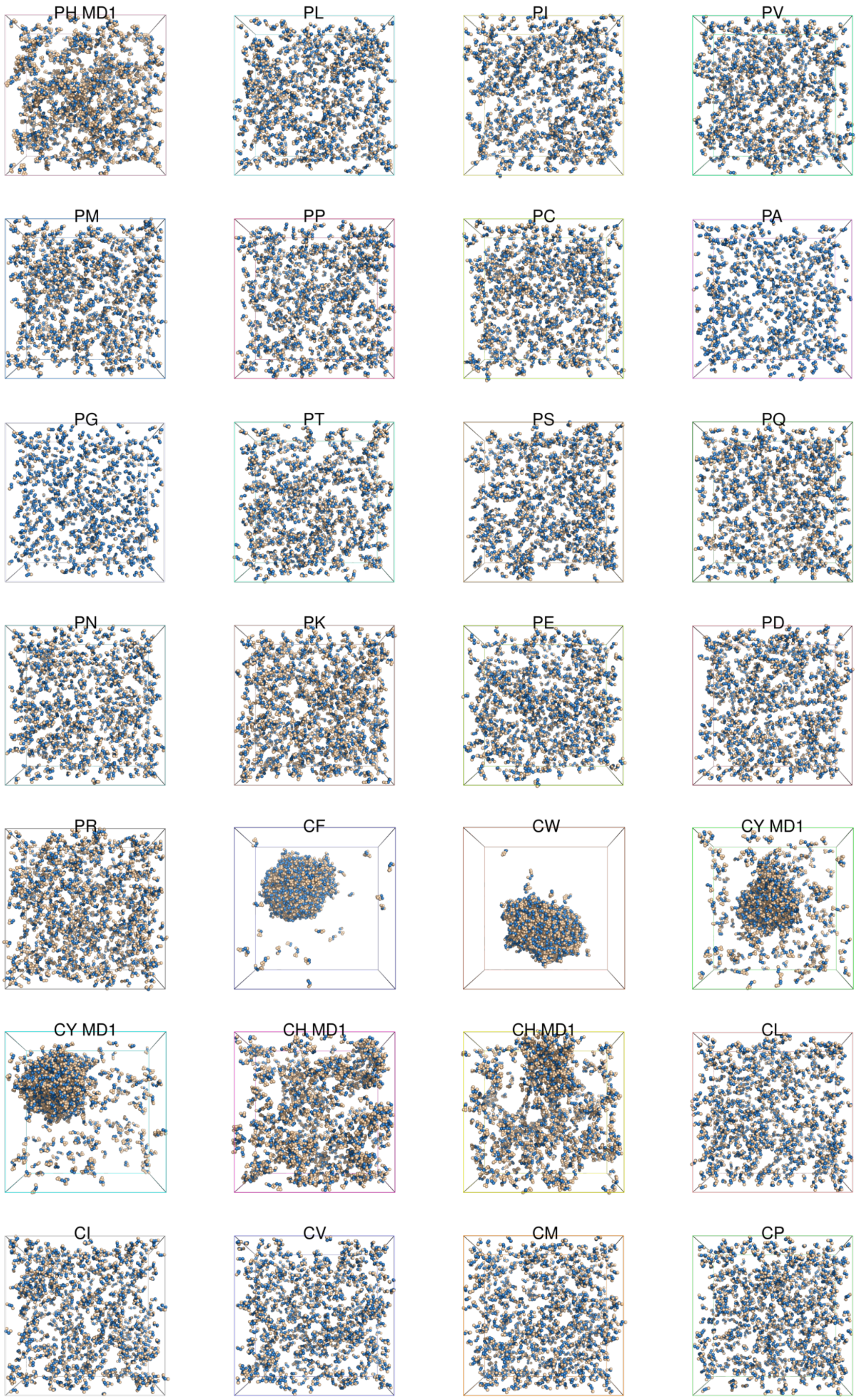
PH MD1
PL
PI
PV
PM
PP
PC
PA
PG
PT
PS
PQ
PN
PK
PE
PD
PR
CF
CW
CY MD1
CY MD1
CH MD1
CH MD1
CL
CI
CV
CM
CP

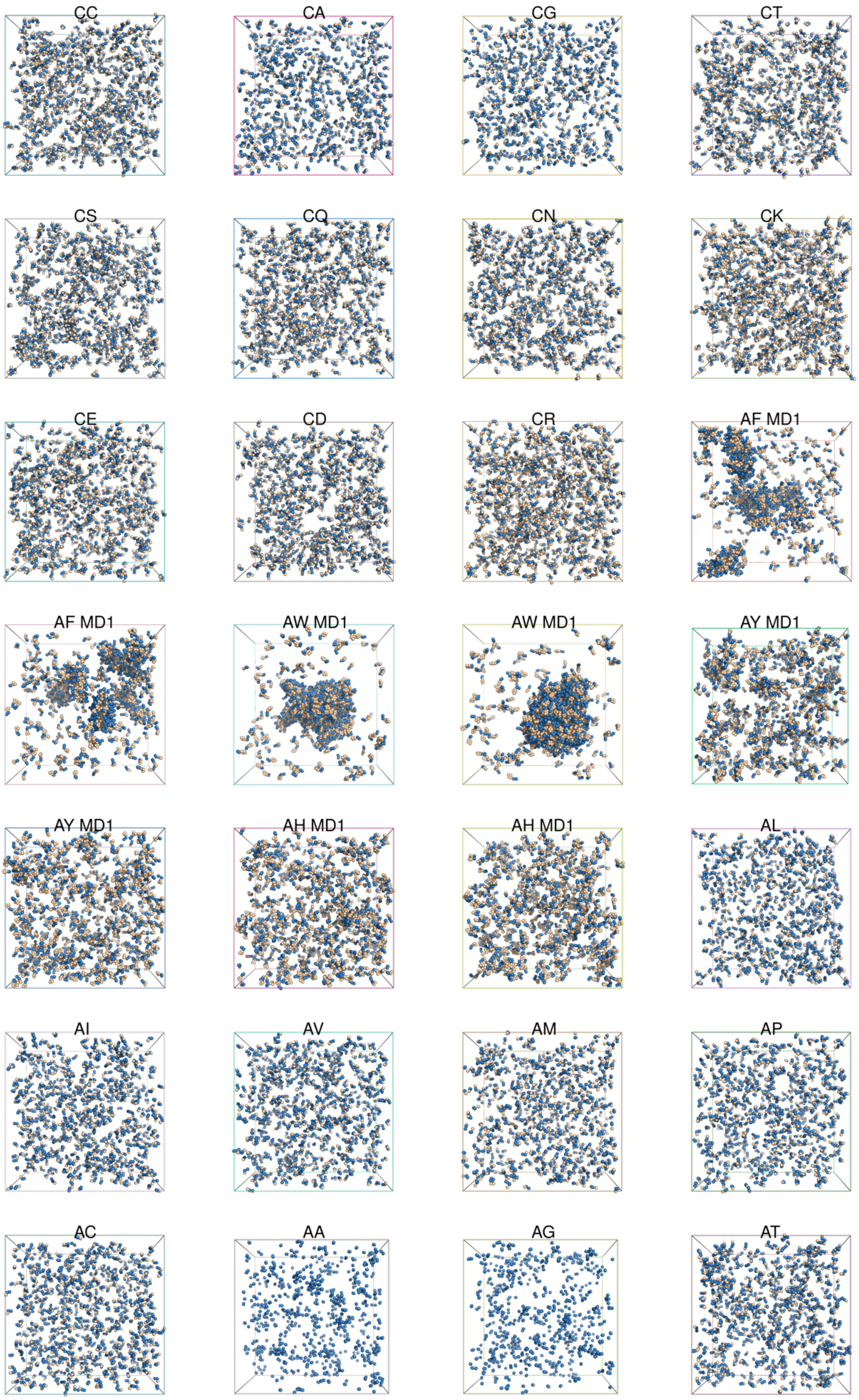
CC
CA
CG
CT
CS
CQ
CN
CK
CE
CD
CR
AF MD1
AF MD1
AW MD1
AW MD1
AY MD1
AY MD1
AH MD1
AH MD1
AL
AI
AV
AM
AP
AC
AA
AG
AT

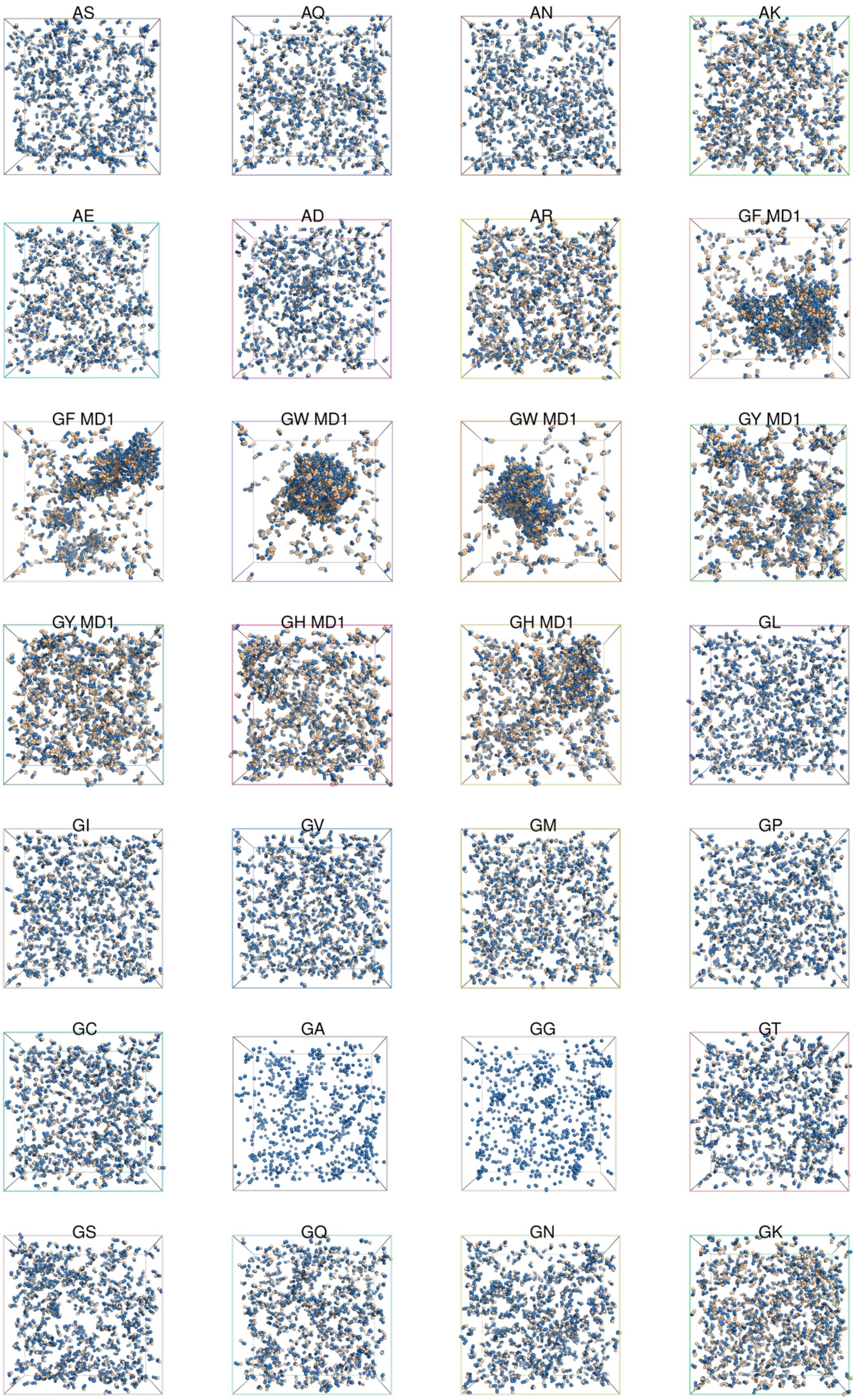
AS
AQ
AN
AK
AE
AD
AR
GF MD1
GF MD1
GW MD1
GW MD1
GY MD1
GY MD1
GH MD1
GH MD1
GL
GI
GV
GM
GP
GC
GA
GG
GT
GS
GQ
GN
GK

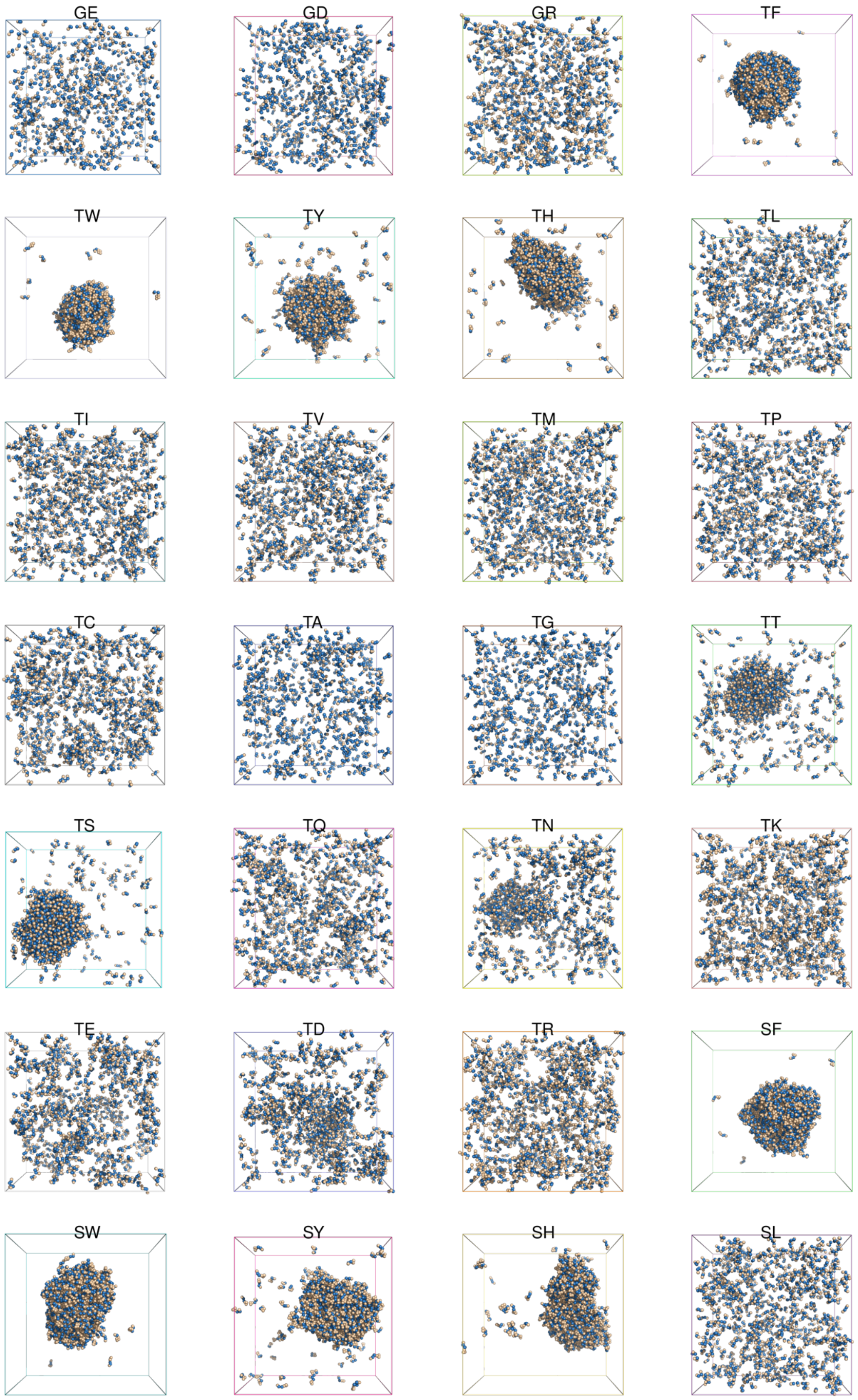
GE
GD
GR
TF
TW
TY
TH
TL
TI
TV
TM
TP
TC
TA
TG
TT
TS
TQ
TN
TK
TE
TD
TR
SF
SW
SY
SH
SL

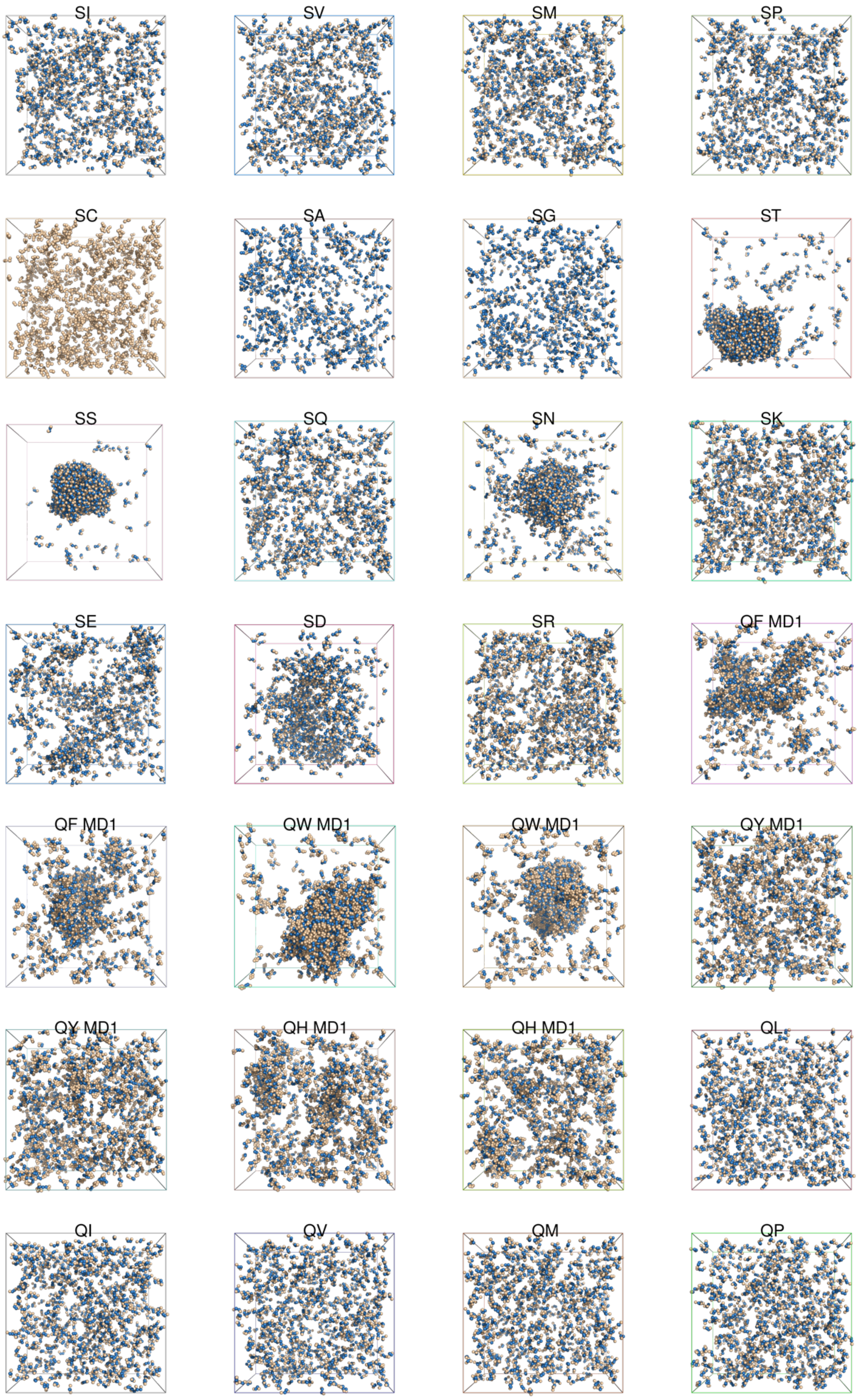
SI
SV
SM
SP
SC
SA
SG
ST
SS
SQ
SN
SK
SE
SD
SR
QF MD1
QF MD1
QW MD1
QW MD1
QY MD1
QY MD1
QH MD1
QH MD1
QL
QI
QV
QM
QP

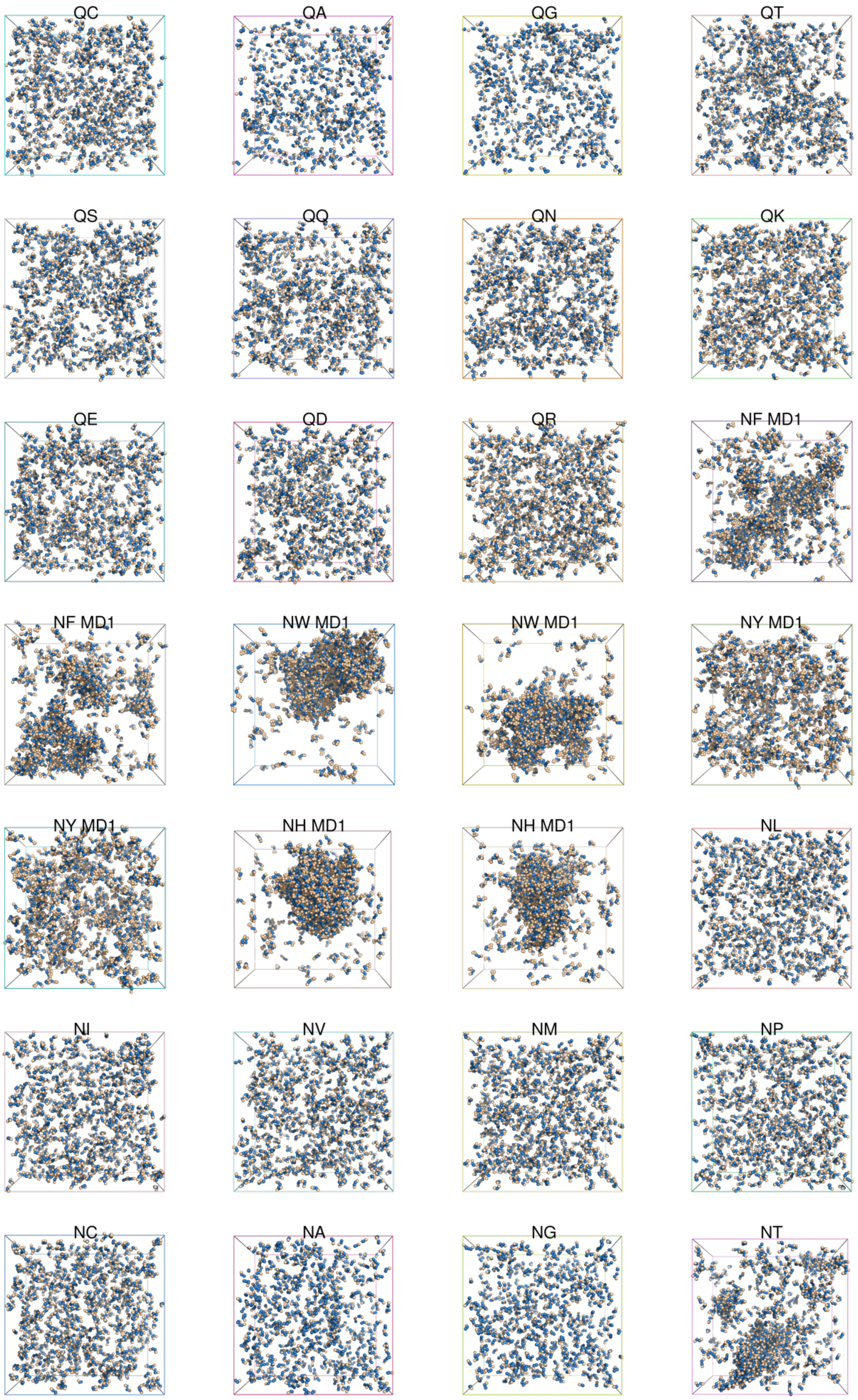
QC
QA
QG
QT
QS
QQ
QN
QK
QE
QD
QR
NF MD1
NF MD1
NW MD1
NW MD1
NY MD1
NY MD1
NH MD1
NH MD1
NL
NI
NV
NM
NP
NC
NA
NG
NT

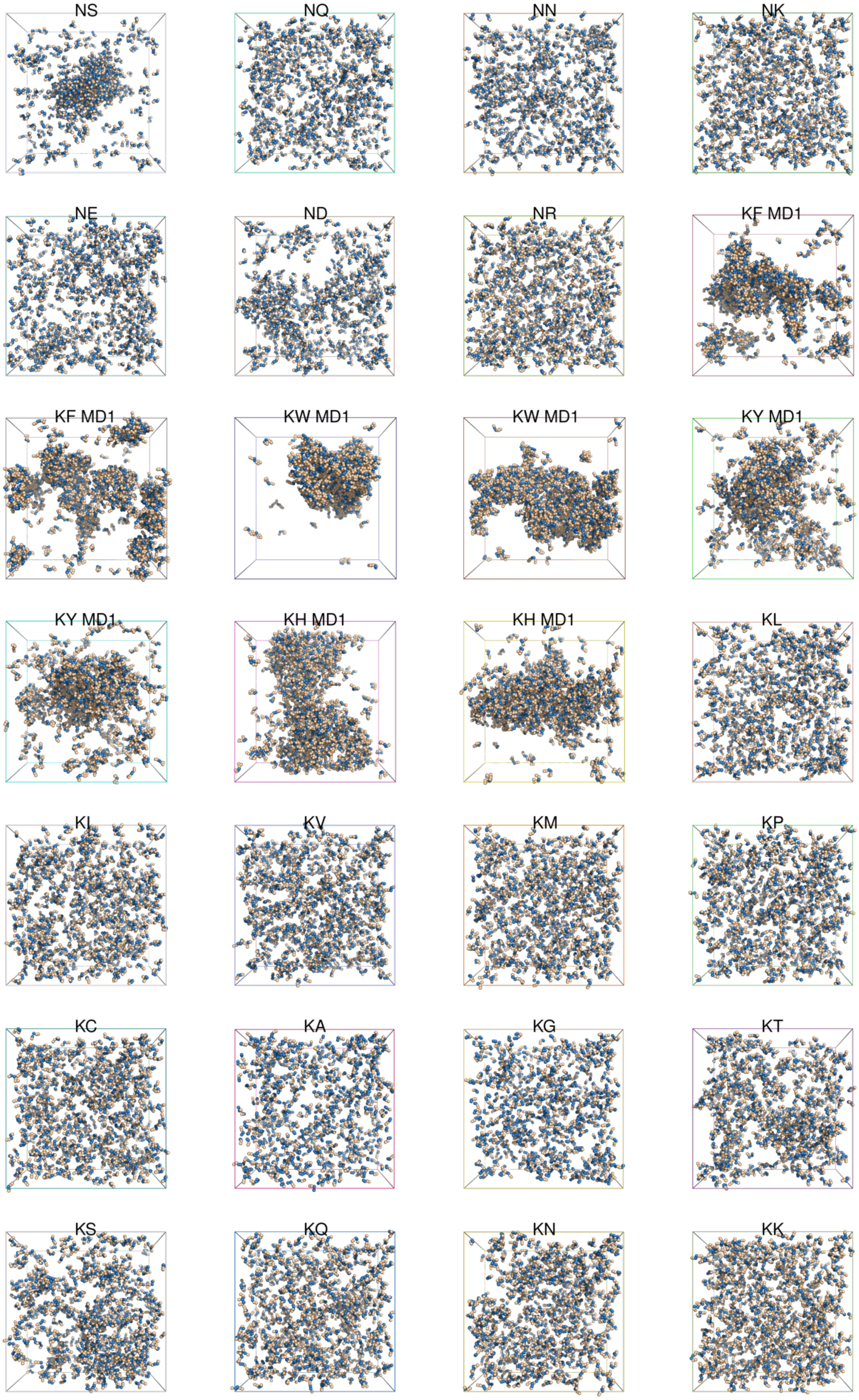
NS
NQ
NN
NK
NE
ND
NR
KF MD1
KF MD1
KW MD1
KW MD1
KY MD1
KY MD1
KH MD1
KH MD1
KL
KI
KV
KM
KP
KC
KA
KG
KT
KS
KQ
KN
KK

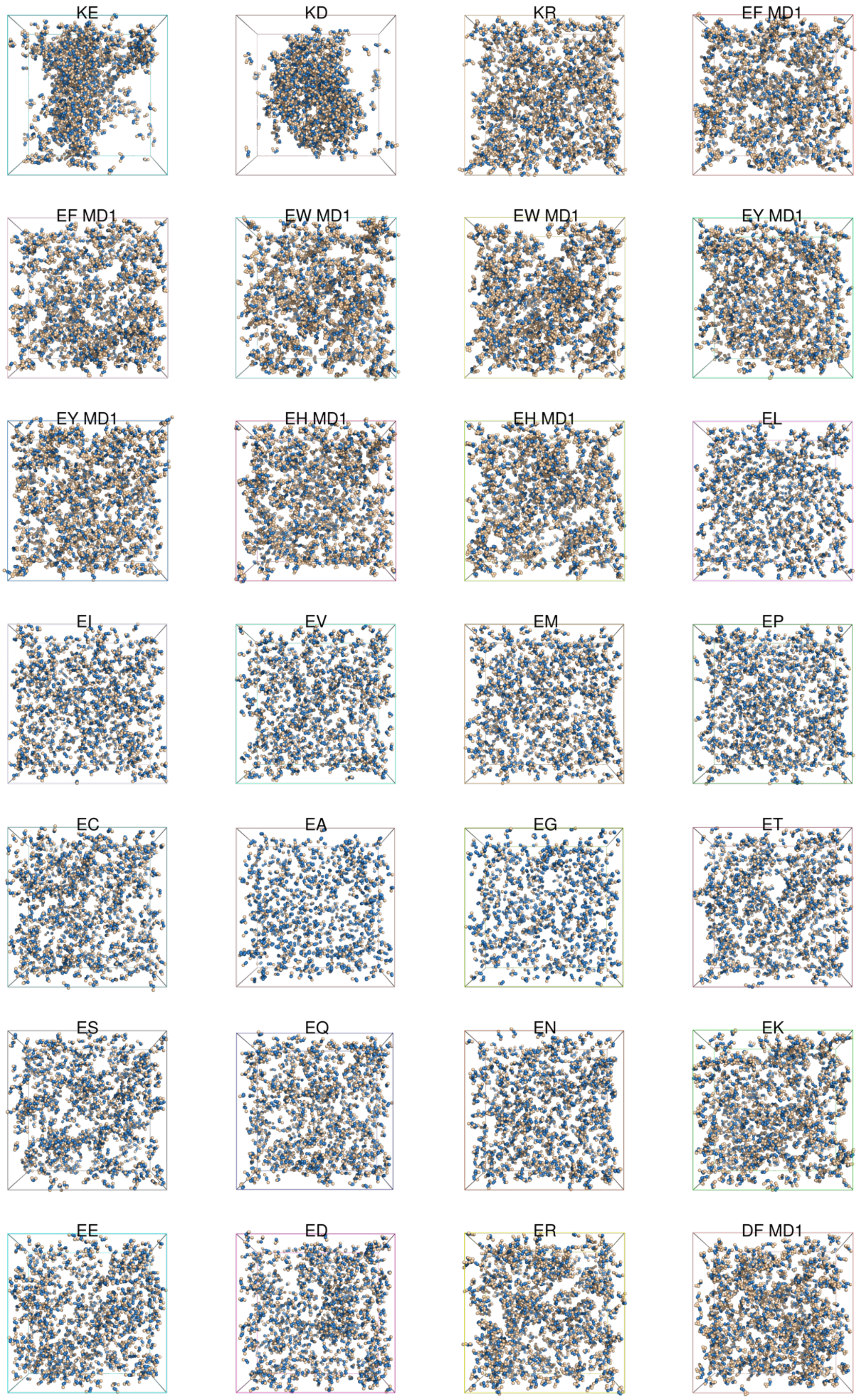
KE
KD
KR
EF MD1
EF MD1
EW MD1
EW MD1
EY MD1
EY MD1
EH MD1
EH MD1
EL
EI
EV
EM
EP
EC
EA
EG
ET
ES
EQ
EN
EK
EE
ED
ER
DF MD1

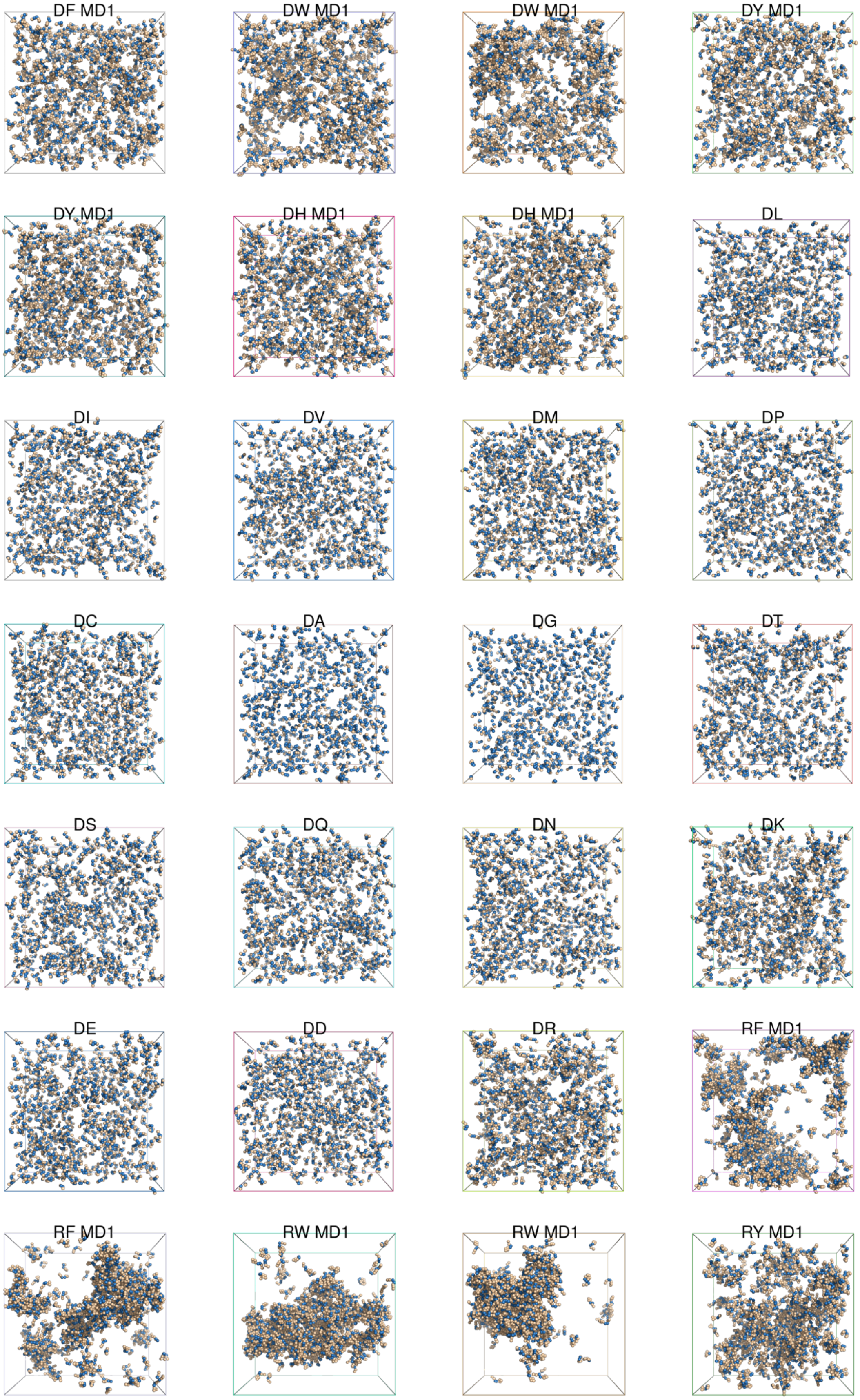
DF MD1
DW MD1
DW MD1
DY MD1
DY MD1
DH MD1
DH MD1
DL
DI
DV
DM
DP
DC
DA
DG
DT
DS
DQ
DN
DK
DE
DD
DR
RF MD1
RF MD1
RW MD1
RW MD1
RY MD1

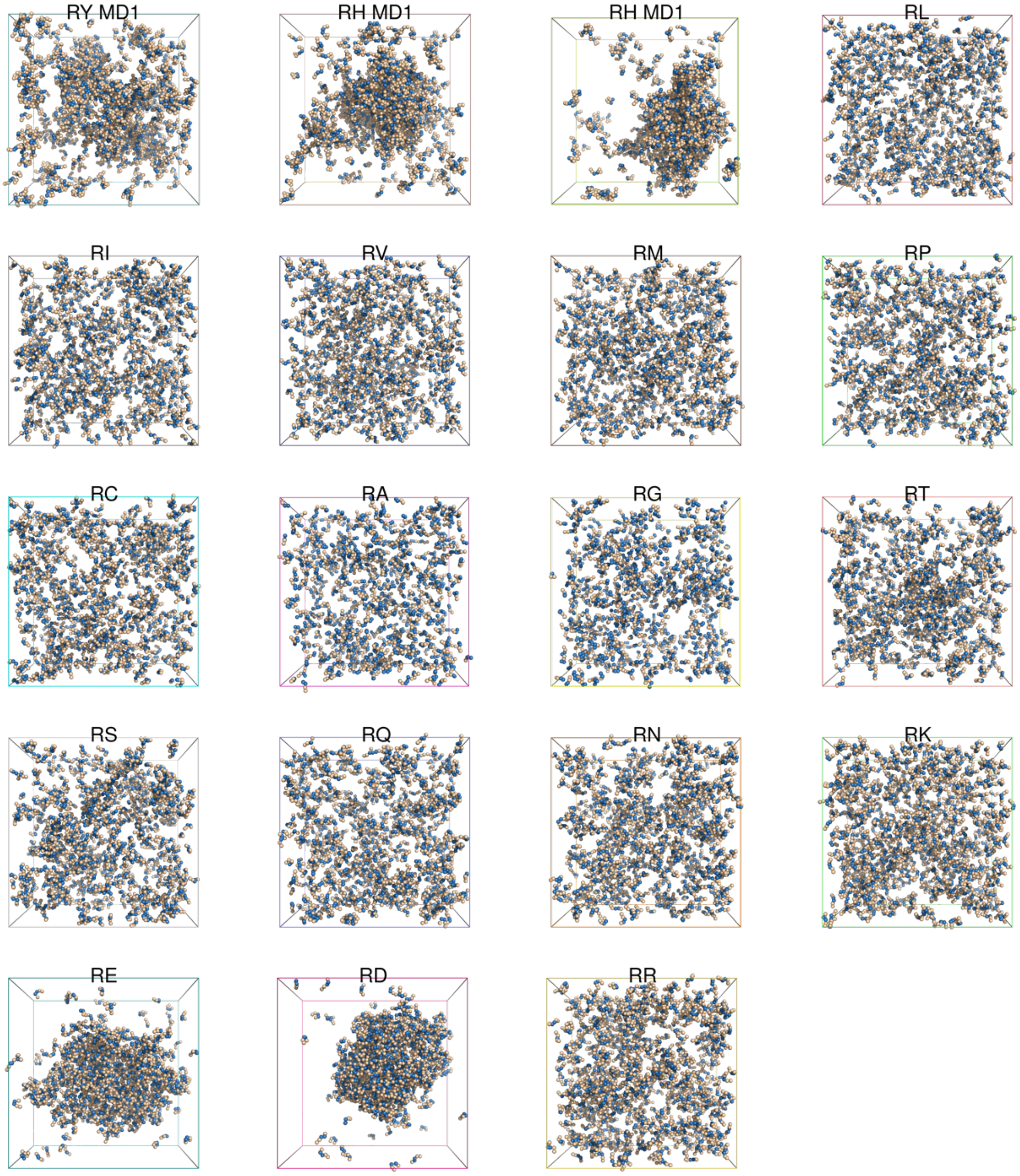
RY MD1
RH MD1
RH MD1
RL
RI
RV
RM
RP
RC
RA
RG
RT
RS
RQ
RN
RK
RE
RD
RR
